# SRPK3 regulates alternative pre-mRNA splicing required for B lymphocyte development and humoral responsiveness

**DOI:** 10.1101/759829

**Authors:** Tessa Arends, J. Matthew Taliaferro, Eric Peterman, Jennifer R. Knapp, Brian P. O’Connor, Raul M. Torres, James R. Hagman

## Abstract

Alternative splicing (AS) of pre-mRNA is a critical component of transcriptional regulation that diversifies the cellular proteome. The Serine-Arginine Protein Kinases (SRPK) initiate early events in AS. Using conditional knockout mice (*cKO*), we demonstrated the importance of the X-linked gene *Srpk3* in B lymphocyte development and in response to immunization in vivo. Significantly decreased numbers of immature and mature B cells were observed in *Srpk3-cKO* BM relative to wild-type (*WT*). Immunization of *Srpk3-cKO* mice with a T lymphocyte-independent type-2 antigen elicited greatly reduced amounts of specific IgG3. *Srpk3* deletion resulted in hundreds of differentially spliced mRNAs in B cells, including mRNAs encoding proteins associated with signaling pathways and mitochondrial function. Several alternative splicing outcomes in *Srpk3-cKO* cells are due to altered splicing regulation of SR proteins. We conclude that *Srpk3* is an immunomodulatory kinase that controls humoral immunity via its regulation of pre-mRNA splicing, antibody production, and metabolism in B cells.

**One Sentence Summary:** SRPK3 regulates alternative splicing of pre-mRNA that is crucial for B cell development, activation and antibody responses.

## Introduction

Co-transcriptional mechanisms of gene regulation are emerging as important contributors to immune cell differentiation and function. One such mechanism is alternative pre-mRNA splicing (AS), which increases informational diversity, functional capacity, and the efficiency of protein expression (*1*). Recent studies have confirmed that AS influences immunity and tolerance through its roles in lymphocyte homeostasis (*2*). Additionally, B cell differentiation, activation, and maturation to antigen secreting cells (ASCs) are dependent on unique splicing patterns that modulate B cell responsiveness and function (*3, 4*). Therefore, understanding how pre-mRNA splicing is regulated in these cells is of critical importance.

The functional spliceosome is a dynamic machine comprised of approximately 20 protein and RNA components that regulate splice site selection in pre-mRNAs concurrently with transcription (*5, 6*). Upstream regulators of splicing include the Serine-Arginine Protein Kinase (SRPK) family, a group of highly conserved serine-threonine kinases that phosphorylate Serine-Arginine repeat (SR)-rich proteins in response to extracellular signals (*7–10*). SR proteins include splicing factors (SRSF1-12) that promote splice site selection by bridging between initial splice site selection in the pre-spliceosome and assembly of the mature spliceosome.

In mammals, SRPK proteins (SRPK1 and SRPK2) have been extensively studied as regulators of AS and mRNA maturation (*11–13*), transduction of growth signaling (*14*), chromatin reorganization (*15*), cell cycle (*7*) and metabolic signaling (*16*). However, the function of SRPK3 has not been elucidated. SRPK3 was identified first as a transcriptional target of Myocyte Enhancer Factor 2C (MEF2C) in skeletal muscle and the heart (*17*). Later studies identified the X-linked *Srpk3* gene as a target of MEF2C and the B lineage master regulator PAX5 in B cells (*18–20*). Notably, expression patterns of *Srpk3* in BCR^+^ B cells of the bone marrow (BM) and periphery suggest that the kinase is important for B cell signaling and antibody responses.

Here, we used B cell-specific conditionally deleted *Srpk3* (*Srpk3-cKO*) mice to demonstrate that SRPK3 is essential for efficient development of immature and mature B cells in the BM. Decreased generation of these cells correlates with extensive changes in AS events. *Srpk3-cKO* mice secrete reduced levels of antigen-specific IgG3 in response to the T lymphocyte-independent type 2 (TI-2) antigen 4-hydroxy-3-nitrophenylacetyl-Ficoll (NP-Ficoll). We confirmed that this phenotype is due to the impairment of marginal zone (MZ) B cells, which also exhibit changes in AS in the absence of SRPK3. Overall, loss of SRPK3 profoundly affects B cell development and cellular metabolism including mitochondrial biogenesis and function necessary for normal immune responses of B cells.

## Results

### Deletion of Srpk3 in vivo results in decreased immature and mature BM B cells

To assess the physiological roles of SRPK3 in B cell development *in vivo*, we bred *Srpk3*-floxed female (*Srpk3^fl/fl^*) or male (*Srpk3^fl/y^*) mice with *CD79aCre^Tg/+^* mice that express Cre recombinase in early B cell progenitors (*21, 22*). Deletion of *Srpk3* floxed alleles results in loss of exons 6-8, which encode functional kinase residues (Fig. S1A). In resulting *Srpk3* conditional knockout (*Srpk3^cKO/cKO^* females or *Srpk3^cKO/y^* males; collectively *Srpk3-cKO*) mice, Cre-mediated deletion initiates in early pre-pro B cells prior to expression of *Srpk3* (Fig. S1 B-D).

To characterize phenotypic differences in BM B cell progenitors lacking SRPK3, we used flow cytometry to determine relative frequencies of cells in BM of wild-type (*WT*) and *Srpk3*-*cKO* male and female mice (*23*). In *Srpk3-cKO* BM, early stages of B cell development are unaffected, while later stage immature and mature B cells are decreased (Fig. 1, A-E). In contrast, we observed no differences between splenic B cell populations of *WT* and *Srpk3-cKO* mice (Fig. S2, A-E).

**Fig. 1.**
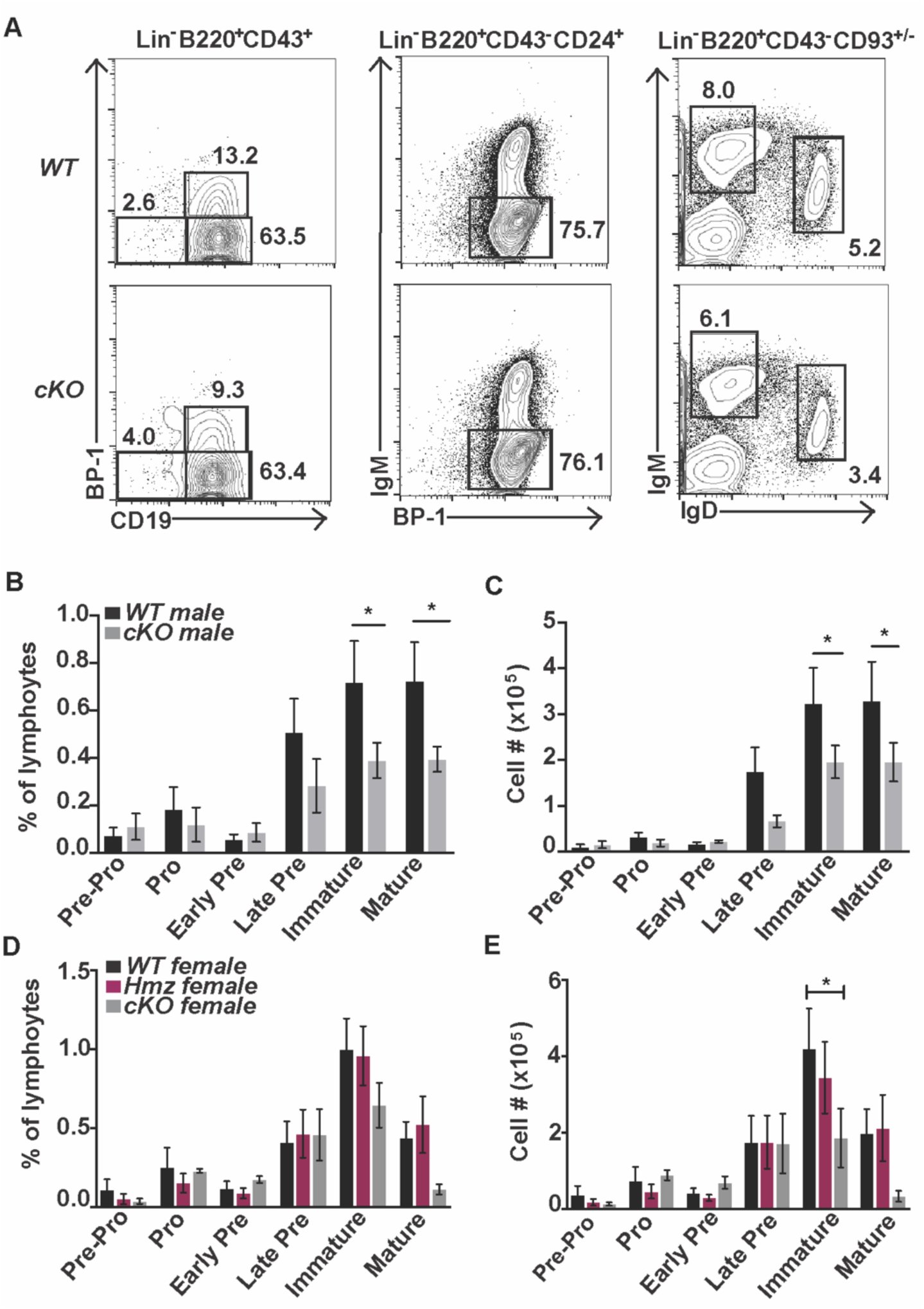
Decreased immature and mature BM B cell populations in *Srpk3-cKO* mice. Quantitation of BM B cell subsets in mice. (**A**) Phenotype, (**B** and **D**) frequencies and (**C** and **E**) cell numbers of various BM B cell populations in male *WT* (n=7-9) or *Srpk3-cKO* (n=7-9) mice (**B** and **C**) and female *WT* (n=3-7), *Srpk3-hemizygous* (*Hmz*)(n= 6-8) and *Srpk3-cKO null* (n=3-5) (**D** and **E**) mice. In panel A numbers indicate percent of phenotypic fraction. Asterisks indicate statistical significance compared to *WT* littermate controls using 2-way ANOVA. *P<0.05. Graphs represent arithmetic mean ± SEM. All data are representative of at least three independent experiments.

The decrease in immature BM B cell numbers in *Srpk3-cKO* mice could be due to impairment of Ig light chain gene rearrangements. Using flow cytometry, we observed that *Srpk3-cKO* immature and mature B cells have increased ratios of κ/λ light chains relative to *WT* cells (Fig. 2, A-D). *S*plenic B cells also show this trend (Fig. S3, A and B). No significant differences were detected between frequencies of intracellular κ^+^ cells or κ mean fluorescence intensity (MFI) in *Srpk3-cKO* versus *WT* pre-B cells (Fig. S4, A and B). Thus, SRPK3 is important for the generation of normal BCR repertoires in mice.

**Fig. 2.**
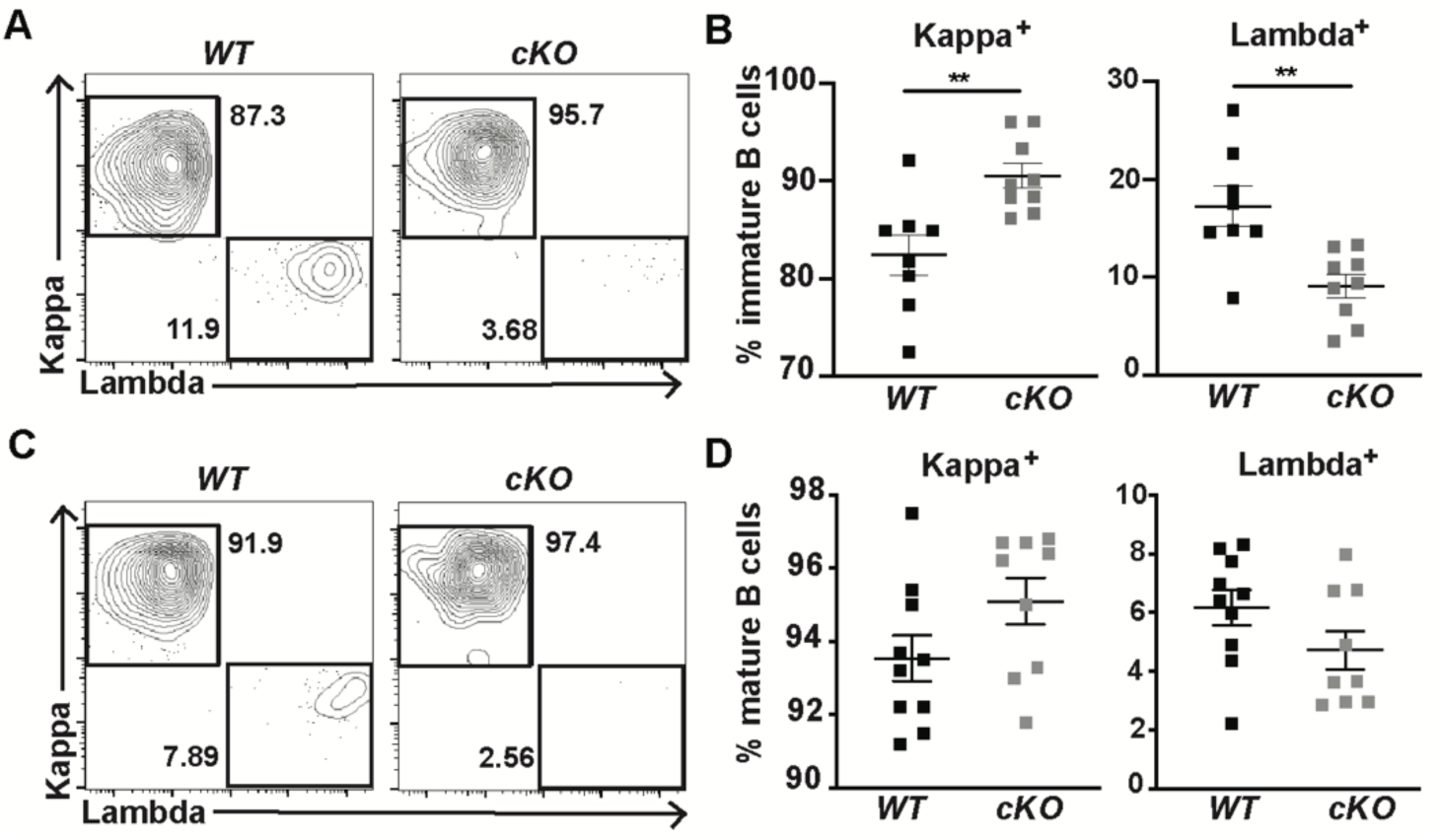
Skewed κ and λ light chain expression on immature BM B cells in *Srpk3-cKO* mice. (**A**) Phenotype and (**B**) frequencies of κ and λ light chain expression on immature BM B cells (CD19^+^Lin^-^B220^+^IgM^+^) from *WT* (n=8) and *Srpk3-cKO* (n=9) male mice. (**C**) Phenotype and (**D**) frequencies of κ and λ light chain expression on mature BM B cells (CD19^+^Lin^-^B220^hi^IgM^+^) from *WT* (n=10) and *Srpk3-cKO* (n=9) male mice. Each dot represents an individual mouse. Asterisks indicate statistical significance compared to *WT* littermate controls using Mann-Whitney Rank test. **P<0.005. Graphs represent arithmetic mean ± SEM. All data are representative of at least three independent experiments.

### SRPK3 regulates patterns of AS in immature and mature BM B cells

*Srpk3* expression is highest in immature B cells during B lymphopoiesis (*24*). Therefore, we hypothesized that SRPK3 regulates splicing of pre-mRNAs necessary to generate immature B cells, and/or for their selection in the BM. For comparison, we analyzed mRNAs isolated from mature BM B cells (*25*). We performed RNA-seq to determine genome-wide changes in transcript frequencies and patterns of AS. Purity of sorted populations was assessed (Fig. S5A) and principal component analysis (PCA) of RNA-seq was performed to determine relationships between biological replicates between populations (Fig. S5B). Differential gene expression (DGE) analysis identified only 5 differentially expressed genes between *WT* and *Srpk3*-*cKO* immature B cells, including *Cd79a* (due to the *Cre* knock-in allele) (≥ 1.5 fold; *P_adj_* < 0.05). Similarly, only 18 genes were differentially expressed between *WT* and *Srpk3*-*cKO* mature B cells, including *Col5a3*, which encodes α chains of fibrillar collagens that regulate metabolism in mice (*26*), and *Fms-tyrosine kinase 3 (Flt3),* which is essential for B cell maturation (*27*) (≥ 1.5 fold, P_adj_ < 0.05) (Data file S1). Analysis of AS events revealed robust changes in splicing patterns between mRNAs of *WT* and *Srpk3*-*cKO* immature and mature BM B cells. We detected 290 significant AS events in immature B cells and 620 significant AS events in mature B cells (ΔΨ ≥ 0.1, FDR < 0.05) (Fig. 3A; Fig. S6A; Data file S2). In both immature and mature B cells, skipped exons (SE) comprised the majority of AS events, while differences in the inclusion of mutually exclusive exons (MXE) were the second most frequent type of AS. Other types of events, including alternative 3’ splice sites (A3SS), alternative 5’ splice sites (A5SS) and retained introns (RI) were detected at lower frequencies. When displayed in a heat map that separates *WT* from *Srpk3-cKO* AS events (Fig. 3A; Fig. S6A), it is apparent that different types of AS are detected both in the absence or presence of SRPK3. Next, we used Ingenuity Pathway Analysis (IPA) to identify pathways affected by AS in immature and mature B cells. Cellular pathways affected in *Srpk3*-*cKO* immature B cells include Ataxia-Telangiectasia Mutated (ATM) signaling, DNA double-strand break (DSB) repair, DNA damage response (DDR) pathways and cell cycle damage checkpoints, Hypoxia signaling and AMPK (Fig. 3B). Additionally, IPA indicated that critical cellular pathways including ERK5, AMPK, mTOR and TGF-β signaling are affected by the loss of SRPK3 in mature B cells (Fig. S6B). In support of RNA-seq data we validated AS of genes that are strongly affected by the loss of SRPK3. AS of transcripts of *Tyrosine kinase 2* (*Tyk2)*, a component of type I and type III interferon signaling pathways that regulates mitochondrial function (*28, 29*), and *NIMA related kinase 4 (Nek4),* which regulates DDR (*30*), were confirmed (Fig.3, C and D). We hypothesize that the combined dysregulation of crucial genes contributes to the decreased frequencies of immature and mature B cells in *Srpk3*-*cKO* mice.

**Fig. 3.**
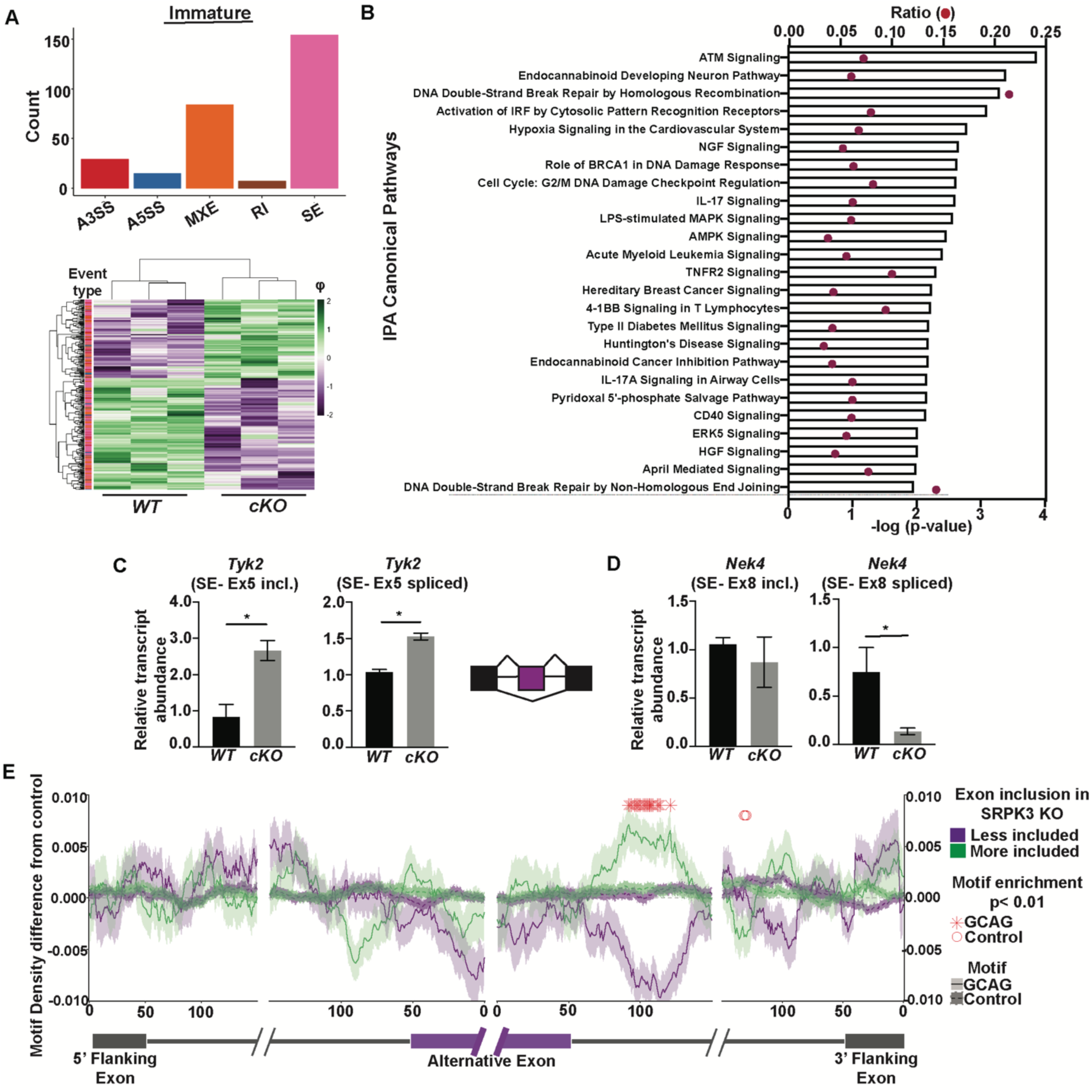
SRPK3 regulates alternative splicing patterns in immature BM B cells. (**A**) Significantly altered splicing event types in purified immature BM B cells between *WT* (n=3) and *Srpk3-cKO* (n=3) male mice. AS event frequencies are displayed for alternative 3’ splice sites (A3SS), alternative 5’ splice sites (A5SS), mutually exclusive exons (MXE), retained introns (RI), and skipped exons (SE). Heatmaps show ψ values. Color (purple to green) represents z-score of ψ values across rows. Δψ ≥ 0.1, FDR < 0.05. No overlap in ψ between replicates across conditions. (**B**) IPA enrichment for the top 30 most significantly enriched canonical pathways in the immature BM B cell RNA-seq splicing dataset. (**C**) Validation (qRT-PCR) of *Tyk2* differential splicing. (**D**) Validation (qRT-PCR) of *Nek4* differential splicing. (**E**) Motif density surrounding SPRK3-sensitive exons. GCAG motif densities were calculated in the sequences surrounding exons that were more included or less included in the *Srpk3-cKO* relative to *WT*. Densities were also calculated for unaffected control exons. The difference in densities between affected and control exons is plotted on the y-axis, with the shaded area representing the standard deviation of the difference. As a further control, the average motif density for a set of control motifs was calculated (dotted line). These motifs were all 4-mers that contained the same GC and CpG content as GCAG. P-values for the enrichments were calculated using a Mann-Whitney U test comparing densities in affected and control exons. Positions were highlighted as significantly enriched (red asterix or circle) if P<0.01.

The most direct way that SPRK3 can affect splicing is through its phosphorylation of SR proteins (*31*). Analysis of the preferred RNA motifs and activities for many RNA binding proteins indicated that SR proteins generally bind the sequence ‘GCAG’ in pre-mRNAs (*32*). Binding of SR proteins to these motifs downstream of alternative exons can inhibit their inclusion in spliced transcripts. Therefore, we expect to observe an enrichment of GCAG motifs downstream of alternative exons that display increased inclusion in *Srpk3-cKO* B cells. We calculated GCAG motif density in regions surrounding alternative exons that were more or less included in *Srpk3-cKO* cells compared to *WT* (Fig. 3E; Fig. S6C). Motif densities in sequences surrounding these exons were compared to densities around control exons that were unaffected in *Srpk3-cKO* cells. As a further control, we also calculated densities for control motifs that had the same GC and CpG contents as GCAG.

In immature B cells, we observed that exons that were more included in *Srpk3-cKO* cells had a significant enrichment of GCAG motifs downstream of the alternative exon compared to exons whose splicing was unaffected (Fig. 3E). Conversely, we observed significantly lower frequencies of GCAG motifs following exons whose inclusion was decreased in *Srpk3-cKO* cells. These observations are consistent with splicing regulatory activities of SRPK3 and SR proteins in immature B cells. We did not observe a similar motif enrichment in the mature B cell data, perhaps due to different regulatory roles of SRPK3 in mature B cells (Fig. S6C).

### SRPK3 is essential for TI-2 immune responses in vivo

Genetic database searches [MyGene2, (*33*); MARRVEL, (*34*)] yielded valuable information regarding known SRPK3 mutations and phenotypes, including immunodeficiency, associated with genetic variants in human patients. To confirm that SRPK3 deficiency attenuates immune function, we first investigated basal immunoglobulin levels in naïve mice. IgG1 serum levels were significantly elevated in naïve *Srpk3*-*cKO* mice, whereas IgG3 serum levels trended lower (Fig. S7A). To determine the impact of SRPK3-deficiency on immune responses *in vivo*, we immunized mice with NP-Lipopolysaccharide (LPS), a model TI-1 antigen, or NP-Ficoll, a model TI-2 antigen, and measured NP-specific IgM and IgG3 antibody responses. Similar levels of NP-specific IgM and IgG3 were detected between *Srpk3*-*cKO* and *WT* mice in response to NP-LPS (over 28 days post-immunization; Fig. S8A). Anti-NP IgM levels were unaffected between *Srpk3*-*cKO* and *WT* mice after immunization with NP-Ficoll. However, *Srpk3*-*cKO* mice produced significantly reduced levels of NP-specific IgG3 over three weeks post-immunization (Fig. 4, A and B). Immunization was also performed with the model T lymphocyte-dependent (TD) antigen NP-chicken-gammaglobulin (NP-CGG), and NP-specific IgM and IgG1 titers were measured. Anti-NP IgM titers were decreased in *Srpk3*-*cKO* mice after initial challenge (Fig. S8B). Additionally, *Srpk3*-*cKO* mice generated increased levels of NP-specific IgG1 antibodies after primary immunization but failed to increase production after secondary immunization (Fig. S8B).

**Fig. 4.**
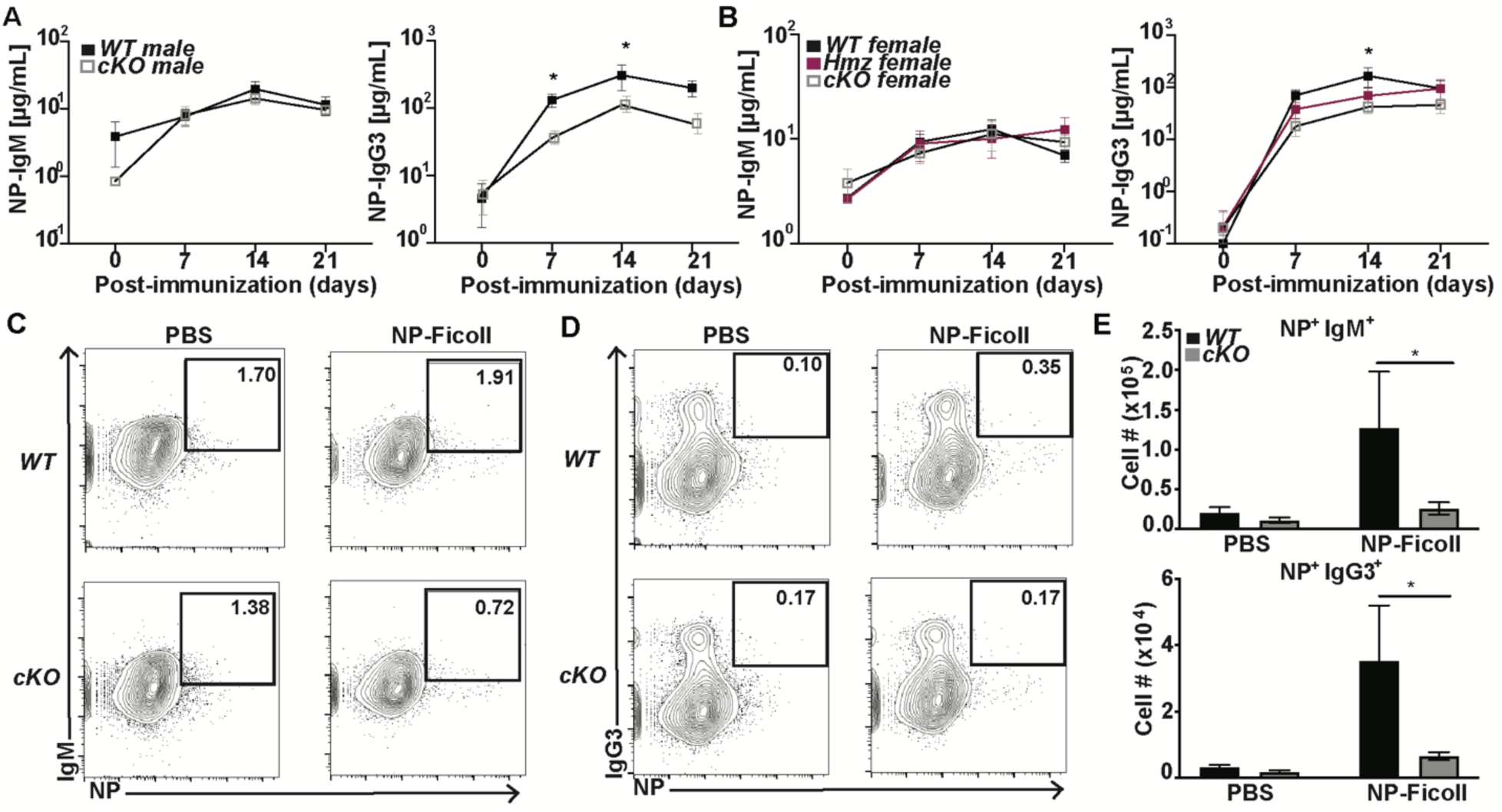
SRPK3 is indispensable for T lymphocyte-independent type-2 humoral immune responses in vivo. (**A**) NP-specific ELISA titers of IgM and IgG3 post-immunization with NP-Ficoll in male *WT* (n=7-8) and *Srpk3-cKO* (n=7) mice. (**B**) NP-specific ELISA titers of IgM and IgG3 post-immunization with NP-Ficoll in female *WT* (n=6-8), *Srpk3-Hmz* (n= 6-7) and *Srpk3*-*cKO* (n=4-6) mice. All points at d14 are significant (*P<0.05) (**C** and **D**) Flow cytometry and (**E**) cell numbers of NP-specific IgM or IgG3 splenic B cells (CD19^+^Lin^-^) in control (PBS) or immunized (NP-Ficoll) male *WT* (n=5) and *Srpk3-cKO* (n=5-7) mice. Mice were harvested 4 days post-immunization for analysis. Asterisks indicate statistical significance compared to *WT* littermate controls using 2-way ANOVA. *P<0.05. Statistical significance is between *WT* and *Hmz* females, as well as, *WT* and *Srpk3-cKO* females in (**B**). Graphs represent arithmetic mean ± SEM. All data are representative of at least three independent experiments.

We hypothesized that the dysregulation of TI-2 antibody responses is due to defective generation or proliferation of NP-specific B cells in *Srpk3*-*cKO* mice. Indeed, we observed decreased frequencies and overall cell numbers of NP-specific IgM^+^ and IgG3^+^ B cells in *Srpk3*-*cKO* mice 4 days after immunization with NP-Ficoll (Fig. 4, C-E). These data suggest that either NP-specific SRPK3-deficient B cells do not proliferate to the same extent as WT B cells in response to immunization or have fewer NP-specific B cells prior to immunization. We also considered whether reductions in NP-specific IgG3^+^ B cells are due to defective class switch recombination (CSR) in response to NP-Ficoll immunization (Fig. S9A). Relative expression of transcripts associated with CSR indicated no significant differences between *WT* and *Srpk3*-*cKO* spleen B cells.

### SRPK3 is required for MZ B cell responses to TI-2 antigens

Given that MZ B cells have been reported to be centrally important for TI-2 antibody responses (*35*), we hypothesized that the inability of *Srpk3-cKO* mice to generate robust immune responses to NP-Ficoll is due, in part, to an intrinsic defect in MZ B cells. Therefore, we purified naïve MZ B cells from *Srpk3-cKO* and *WT* (all male) mice and stimulated them in vitro with dextran-conjugated anti-IgD antibody (α-δ-dex), which elicits TI-2 type BCR signaling, together with interleukin-5 (IL-5) and interferon-gamma (IFNγ) to mimic the response to soluble TI-2 antigens in vivo (*36, 37*). Initially, we measured IgM and IgG3 antibody secretion after 6 days in culture. *Srpk3-cKO* MZ B cells produced significantly lower titers of both IgM and IgG3 compared to *WT* cells (Fig. 5, A and B). These data suggest that SRPK3 is crucial for MZ B cell responses to antigens *in vitro*. We next quantitated surface expression of BCR and cytokine receptors using flow cytometry. *Srpk3-cKO* naïve mice had significantly reduced IL-5Rα^+^ MZ B cells (Fig. S10, A and B). However, we found no relative differences between IgD or IFNγ-receptor (IFNGR1) expression on MZ B cells from *Srpk3-cKO* versus *WT* unimmunized mice (Fig. S10, C-F). We next determined whether defective antibody production was due to failure of MZ B cells to proliferate, differentiate to ASCs, or undergo CSR. *WT* and *Srpk3-cKO* MZ B cell numbers increased similarly in response to activation and unstimulated cells were detected in comparable numbers after 3 days in vitro (Fig. S11A). Thus, *Srpk3-cKO* MZ B cells proliferate normally in response to BCR stimulation and displayed similar in vitro half-life as WT. Next, cells were harvested and analyzed for differentiation to ASCs using CD138 and TACI as plasma cell markers (*38, 39*). Both *Srpk3-cKO* and *WT* MZ B cells differentiated into ASCs similarly after 3 days in culture (Fig. S12, A and B). To assess CSR, we analyzed germline and post-switch transcripts in naïve and stimulated cells. We detected no defects in the abilities of *Srpk3-cKO* MZ B cells to undergo CSR in response to activation (Fig. S13A). These data indicate that *Srpk3-cKO* MZ B cells proliferate and differentiate normally but have additional underlying abnormalities that influence the ability of these cells to respond to antigen.

**Fig. 5.**
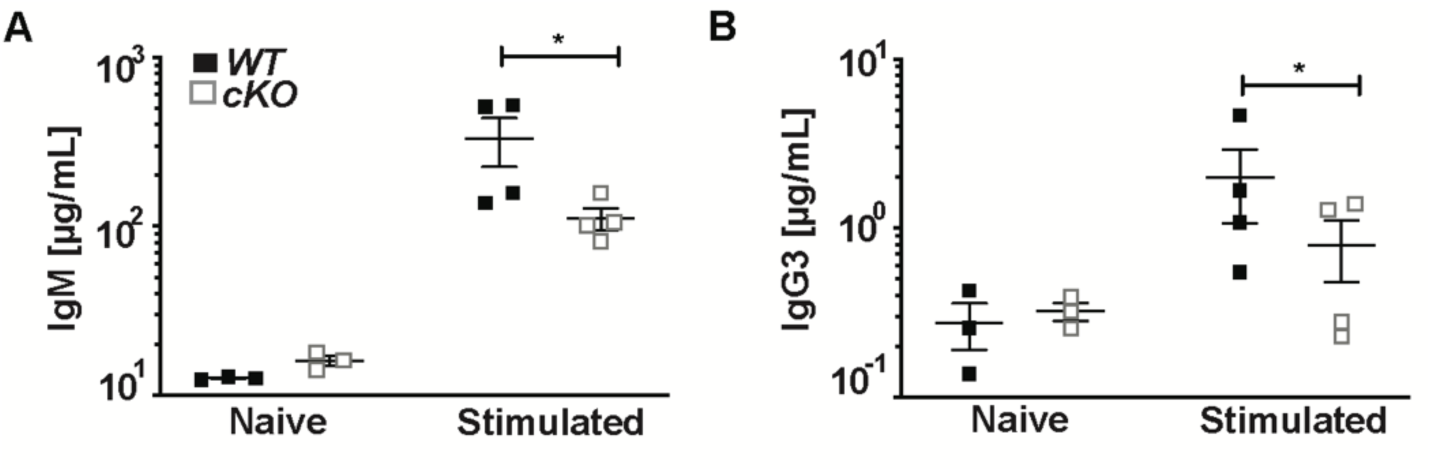
Reduced antibody responses of *Srpk3-cKO* MZ B cells in vitro. (**A**) ELISA detection of IgM antibody in culture supernatants after 6-days of in vitro stimulation (α-δ-dex, IL-5 and IFNγ) of purified MZ B cells from *WT* (n=3-4) and *Srpk3-cKO* (n=3-4) male mice. Each dot represents an individual mouse. (**B**) ELISA detection of IgG3 antibody (same cultures as in **A)**. Asterisks indicate statistical significance compared to *WT* littermate controls using 2-way ANOVA. *P<0.05. Graphs represent arithmetic mean ± SEM. Data are representative of at least three independent experiments.

### SRPK3 modulates alternative splicing patterns in naïve and stimulated MZ B cells

To further delineate mechanisms requiring SRPK3 in MZ B cells, we used RNA-seq to interrogate mRNA sequences in naïve *WT* or *Srpk3-cKO* MZ B cells, without or with stimulation *in vitro* (α-δ-dex, IL-5 and IFNγ) for 8 or 72 hours. Purity of sorted populations was assessed (Fig. S14A) and PCA of RNA-seq was performed to determine relationships between biological replicates (Fig. S14B). DGE analysis yielded no significant differences in overall gene expression between the two genotypes of naïve and stimulated MZ B cells (≥ 1.5 fold, P_adj_ < 0.05) (Data file S3). However, analysis of AS revealed changes in splicing patterns between *WT* and *Srpk3-cKO* MZ B cells, under both naïve and stimulated conditions (Fig. 6, A and B; Fig. S15, A and B). We detected 213 significant alternative splicing events in naïve samples, 38 significant alternative splicing events in 8-hour stimulated samples, and 51 significant alternative splicing events in 72-hour stimulated samples (ΔΨ ≥ 0.1, FDR < 0.05) (Data file S4).

**Fig. 6.**
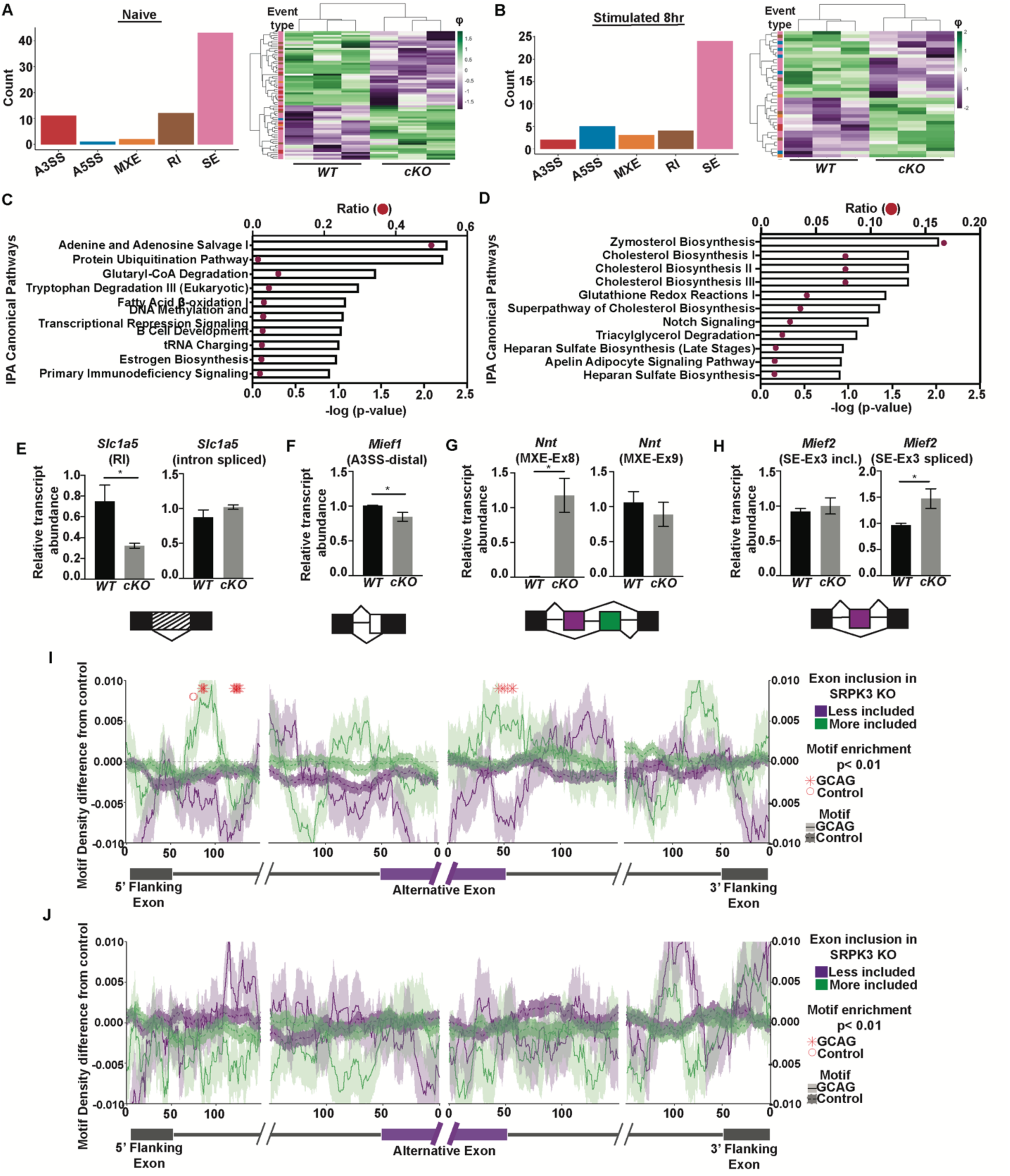
SRPK3 modulates alternative splicing patterns in MZ B cells. (**A**) Significantly altered splicing event types in (**A**) naïve or (**B**) stimulated (8h, as in Fig. 5) MZ B cells purified from *WT* (n=3) and *Srpk3-cKO* (n=3) male mice. AS event frequencies are displayed as in Fig. 3A. (**C**) IPA enrichment for the top 10 most significantly enriched canonical pathways in the naïve MZ B cell RNA-seq splicing dataset. (**D**) IPA enrichment for the top 11 most significantly enriched canonical pathways in the 8-hour stimulated MZ B cell RNA-seq splicing dataset. (**E**) Validation (qRT-PCR) of *Slc1a5* differential splicing. (**F**) Validation (qRT-PCR) of *Mief1* differential splicing. (**G**) Validation (qRT-PCR) of *Nnt* differential splicing. (**H**) Validation (qRT-PCR) of *Mief2* differential splicing. Asterisks indicate statistical significance compared to *WT* littermate controls using unpaired, two-tailed Student’s t-test. *P<0.05. Graphs represent arithmetic mean ± SEM. (**I** and **J**) Motif density surrounding SPRK3-sensitive exons in naïve (**I**) or 8-hour stimulated (**J**) MZ B cells (all details similar to Fig. 3E).

We focused on naïve and 8-hour stimulated MZ B cells, because these conditions captured splicing changes before and after B cell activation. Moreover, RNA-seq confirmed that *Srpk3* gene transcription shuts off between 8- and 72-hours of stimulation (Data file S3). IPA of differentially spliced genes identified several key cellular pathways that were enriched in *WT* versus *Srpk3-cKO* MZ B cells (Fig. 6, C and D). In naïve cells, these pathways included purine metabolism, B cell development, estrogen biosynthesis and primary immunodeficiency signaling (Fig. 6C). In 8-hour stimulated cells, enriched pathways included cholesterol biosynthesis, triacylglycerol degradation and heparan sulfate biosynthesis (Fig. 6D). Because a majority of these pathways were related to cell metabolism, we addressed the hypothesis that differential splicing of genes in *Srpk3-cKO* MZ B cells impairs metabolic function. In naïve cells, validation was obtained for AS of *Solute carrier family 1 member 5 (Slc1a5),* a neutral amino acid transporter of glutamine (*40*), and *Mitochondrial elongation factor 1* (*Mief*1; independently identified as MiD51), which regulates mitochondrial fusion (*41*) (Fig. 6, E and F). In 8-hour stimulated samples, *Nicotinamide nucleotide transhydrogenase* (*Nnt*), an inner mitochondrial membrane protein that contributes to mitochondrial NADPH/NADP^+^ ratios (*42*) and *Mitochondrial elongation factor 2* (*Mief2*; independently identified as MiD49), a mitochondrial outer membrane protein that regulates fusion with MiD51 (Fig. 6, G and H), undergo AS. Splicing differences between *WT* and *Srpk3-cKO* MZ B cells, regardless of stimulation, suggest that SRPK3 fine tunes AS events in MZ B cells.

Similar to mutant immature B cells, we expected to detect the enrichment of SR protein target motifs near exons that were more included in *Srpk3-cKO* MZ B cells. We observed enrichment of GCAG motifs downstream of included exons in *Srpk3-cKO* naïve MZ B cells (Fig. 6I). Conversely, GCAG motifs were less frequent downstream of exons whose inclusion decreased in *Srpk3-cKO* naïve MZ B cells. Control motifs that were matched for GC and CpG content were not similarly enriched. We did not observe a similar motif enrichment in the 8-hour stimulated *Srpk3-cKO* MZ B cells, perhaps due to down-regulation of SRPK3 activity following stimulation (Fig. 6J).

### Mitochondrial dynamics and metabolism are disrupted in splenic B cells that lack SRPK3

Notably, AS of genes that function in mitochondrial maintenance and activity has detrimental effects on cellular metabolism (*43*). Flow cytometric analysis revealed that naïve and stimulated *Srpk3-cKO* MZ B cells each have decreased mitochondrial mass compared to *WT* MZ B cells (this analysis does not consider relative ultrastructural morphology of mitochondria, which may influence the observed decrease in mass) (Fig. 7, A and B). To determine whether the loss of mass is due to a decline in mitochondrial abundance, we stained cells with Mitotracker Red and enumerated relative numbers of mitochondria per cell in naïve and stimulated samples using high resolution microscopy. *Srpk3-cKO* naïve MZ B cells exhibited considerably decreased numbers of mitochondria per cell compared to *WT* cells (Fig. 7C). However, after stimulation mitochondrial frequencies were relatively similar between *WT* and *Srpk3-cKO* cells.

**Fig. 7.**
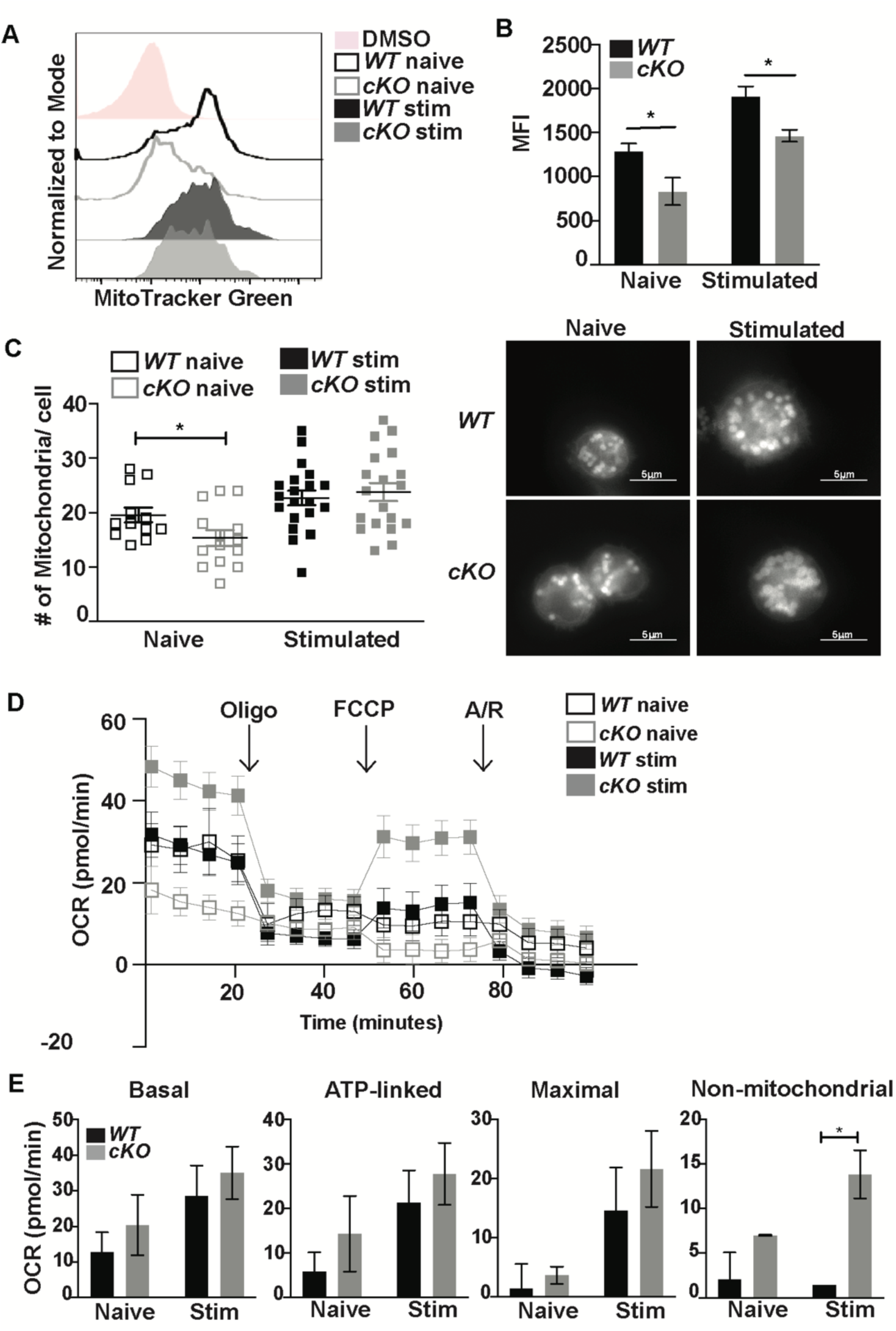
Dysregulation of mitochondrial dynamics and metabolism in *Srpk3-cKO* splenic B cells. (**A**) Flow cytometric analyses and (**B**) MFI of Mitotracker Green in naïve and 24-hour stimulated (α-δ-dex, IL-5 and IFNγ) MZ B cells purified from *WT* (n=3) and *Srpk3-cKO* (n=3) male mice. (**C**) Number of mitochondria per cell in naïve and 24-hour stimulated MZ B cells purified from *WT* (n=2) and *Srpk3-cKO* (n=2) male mice. Each dot represents an individual cell. Representative image of stained mitochondria (Mitotracker Red) in naïve or stimulated (stim) *WT* and *Srpk3-cKO* MZ B cells. 5μm scale. (**D**) OCR measurements between purified naïve or 5-hour stimulated *WT* and *Srpk3-cKO* splenic B cells. (**E**) OCR of basal respiration, ATP-linked respiration, maximal respiration and non-mitochondrial respiration between purified naïve or 5-hour stimulated *WT* and *Srpk3-cKO* splenic B cells. Asterisks indicate statistical significance compared to *WT* littermate controls using either 1-way ANOVA or 2-way ANOVA. *P<0.05. Graphs represent arithmetic mean ± SEM. All data are representative of at least three independent experiments.

We next determined whether AS of genes involved in mitochondrial function affects mitochondrial metabolism. We performed mitochondrial stress tests and measured oxygen consumption rates (OCR) between *WT* and *Srpk3-cKO* MZ B cells. *Srpk3*-deficient naïve and stimulated cells each had increased mitochondrial respiratory capacity when compared to *WT* cells (Fig. 7, D and E). Additionally, the mutant cells had increased non-mitochondrial respiration, which is indicative of decreased mitochondrial function (*44*). Together, these data suggest that mitochondrial function is perturbed in the absence of SRPK3.

## Discussion

Functions of SRPK3 in B cells have remained elusive. Here, we observed striking effects of the deletion of *Srpk3* genes in *cKO* mice. Numbers of B cell progenitors in *Srpk3-cKO* mice are reduced significantly. Antibody responses to TI-2 (and to a lesser degree, TD) antigens were greatly reduced, and the strength of B cell responses to the model TI-2 antigen NP-Ficoll was proportional to the dosage of *WT Srpk3* alleles: *WT*>female hemizygous>null mice, suggesting that *Srpk3* impacts humoral responses in a dosage/sex-dependent manner.

The molecular basis of SRPK3’s regulation of humoral immunity is undoubtedly complex, because deletion of *Srpk3* genes in B cells results in hundreds of different AS events. These results were expected because SRPK family kinases control AS in multiple cell types (reviewed in (*9*). We linked AS in immature, mature and MZ B cells with functions including cell signaling and metabolism. The majority of AS events involved SE and MXE, which can result in alternative protein isoforms or promote instability of mRNA and nonsense-mediated decay. In B cells, we propose that SRPK3 regulates SR proteins that direct exon inclusion and skipping (*45, 46*). We observed the enrichment of RNA sequence motifs that are often bound by SR proteins in sequences surrounding SRPK3-sensitive exons, arguing that at least some of the alternative splicing outcomes in *Srpk3-cKO* cells are due to altered regulation of SR proteins.

Immature B cells represent a key checkpoint in BM B cell development. Reduced numbers of these cells and skewed κ/λ ratios suggest that SRPK3 is involved in generation of the B cell repertoire (*47–49*). Despite clues to the overall functions of SRPK3, the mechanism that reduces BM progenitors in the absence of SRPK3 is unknown. Mutant immature B cells include higher frequencies of Igκ^+^ cells, suggesting that either the *Igλ* loci do not rearrange as efficiently or BCR signaling (and in turn, receptor editing) is impaired in cells lacking SRPK3. Additionally, changes in signaling pathways important for immature B cell development were noted, including the ATM kinase, which orchestrates cellular DSB and DDR responses (*50, 51*) and is critical in regulating V(D)J recombination (*52*). Other pathways included modulators of metabolism and mitochondria. In mature B cells, the ERK5 signaling pathway, which is indispensable for BAFF-induced mature B cell survival and self-reactivity, was affected (*53, 54*), as well as the mechanistic target of rapamycin (mTOR) and transforming growth factor (TGF)-β signaling pathways (*55, 56*). Together, the large number of affected pathways suggests compound effects of mutant proteomes in the absence of SRPK3.

A major conclusion of our studies is that SRPK3 is required for robust humoral responses of B cells *in vivo* and *in vitro*. Although the lack of SRPK3 in peripheral B cell subsets did not affect their numbers in unimmunized mice, antibody production (i.e. IgG3) in response to challenge with the TI-2 antigen NP-Ficoll was depressed in male and female null *Srpk3-cKO* mice. Production of IgG3 in response to TI-2 antigens is a primary role of MZ cells, which are pre-activated to respond to antigens on circulating bacteria in the spleen (*57, 58*). MZ B cells undergo CSR preferentially to the IgG3 isotype, which facilitates discrimination of multivalent antigenic targets (*59, 60*). The multivalent antigen NP-Ficoll primarily elicits a λ^+^-anti-NP antibody response by MZ (and also by B1) B cells. Decreased λ^+^ cells in *Srpk3*-*cKO* mice may, in part, explain the deficit in TI-2 responses of these mice. While we have not examined antibody responses to pathogens, we hypothesize that loss of SRPK3 in mice and humans affects the ability to respond to and clear infection.

Many of SRPK3-regulated transcripts encode proteins with functions in cellular metabolism or in the maintenance or biogenesis of mitochondria. In this regard, recent studies confirmed the importance of metabolic reprogramming as a ‘gatekeeper’ of B lymphocyte development, activation and responses (*61–65*). Loss of SRPK3 in MZ B cells results in differential splicing of candidate gatekeepers *Nnt* and *Slc1a5*. *Nnt* is essential for non-mitochondrial respiration, with important roles in pyruvate metabolism and the Tricarboxylic Acid cycle (TCA) (*66*). In turn, the TCA cycle requires glutamine, which is transported by the *Slc1a5* solute carrier. Additionally, we observed reduced mitochondrial mass in the absence of SRPK3, which could be due to the differential splicing of *Mief1* and *Mief2* transcripts encoding the mitochondrial remodeling factors MiD49/51. MiD49/51 helps recruit essential factors like Drp1 to mitochondria to regulate mitochondrial fission (*41*). Changes in splicing of metabolic genes results in mitochondrial dysfunction, as evidenced by increased OCR in SRPK3-deficient cells. Bioenergetically disrupted mitochondria require higher levels of ATP for maintaining organelle integrity, which increases the basal oxygen consumption rate (*42, 44, 67*). These patterns of high basal OCR and increased glycolysis resemble those in aging cells, which may indicate that the loss of SRPK proteins (*e.g.* SRPK1) contributes to metabolic shifts observed in aging (*68*).

In summary, we propose that SRPK3 is an integral component of splicing pathways in multiple peripheral B cell compartments. Moreover, SRPK3 likely has similar functions in human B cells. A pediatric male patient carrying the mutation I446S in his single *SRPK3* allele is reported to have hypogammaglobulinemia, poor responses to vaccines, and frequent ear/sinus and viral infections in addition to expected muscle weakness (MyGene2; (*33*), while his mother is ‘nearly asymptomatic’. These reports suggest important phenotypes relative to dosage of *SRPK3*. Finally, while SRPK1 and SRPK2 have been studied in human cancers (*69*), roles of SRPK3 and its importance as a biomarker have not been determined. Additional studies are needed to expand our understanding of SRPK3 functions in normal immunity and effects of its dysregulation in human immunodeficiencies and cancer.

## Materials and Methods

### Study Design

The overall goal of these studies was to understand SRPK3’s biological roles in B lymphocyte development and function. This was accomplished by specifically ablating *Srpk3* genes using Cre-Lox technology in B lineage cells *in vivo* and monitoring B lymphocyte development, activation and function as a regulator of pre-mRNA splicing. The observed molecular and cellular changes in B cell function due to the loss of SRPK3 was the basis for conclusions drawn.

### Animals

All mice were maintained on the C57BL/6N genetic background. The *Srpk3-floxed* mice were obtained from KOMP Repository (Davis, CA). The *Cd79a-Cre^Tg/+^* mice (*21*) were kindly provided by John Cambier (University of Colorado, Aurora). Genotyping of *Srpk3* and the *Cd79a-Cre* transgene was performed using tail DNA, PCR and primers as described previously (*70*) (Table S1). All experiments used age-matched male or female test and control littermate mice (6-8 week old). *WT* littermate controls included mice of the following genotypes: *Srpk3^fl/fl^Cd79a^+/+^*, *Srpk3^fl/y^Cd79a^+/+^, Srpk3^WT/WT^Cd79a-Cre^Tg/+^, Srpk3^WT/WT^Cd79a-Cre^+/+^, Srpk3^WT/y^Cd79a-Cre^Tg/+^* and *Srpk3^WT/y^Cd79a-Cre^+/+^*. Mice were bred and housed in the Biological Resources Center at National Jewish Health. All experiments were performed under a protocol (AS2558-04-20) approved by the Institutional Animal Care and Use Committee at National Jewish Health. Further information on the materials and methods used can be found in the Supplementary Materials.

## SUPPLEMENTARY MATERIALS

### Material and Methods

Fig. S1. Efficient deletion of floxed *Srpk3* alleles in BM and splenic B cells.

Fig. S2. Normal splenic B cell development in mice that lack SRPK3.

Fig. S3. κ and λ light chain expression on splenic B cell subsets in *Srpk3-cKO* mice.

Fig. S4. Intracellular κ expression in BM pre-B cells.

Fig. S5. Purity of FACS sorted immature and mature BM B cells and relative variation between RNA-seq replicates.

Fig. S6. SRPK3 regulates alternative splicing patterns in mature BM B cells.

Fig. S7. Elevated IgG1 immunoglobulin levels in *Srpk3-cKO* naive mice.

Fig. S8. T lymphocyte-independent type-1 and T lymphocyte-dependent humoral immune responses in *Srpk3-cKO* mice.

Fig. S9. NP-Ficoll immunized mice are able to undergo class switch recombination.

Fig. S10. Decreased frequencies of IL-5Rα expressing and elevated frequencies of IgD expressing MZ B cells in *Srpk3-cKO* naïve mice.

Fig. S11. *Srpk3-cKO* MZ B cells are capable of proliferating in response to simulation in vitro.

Fig. S12. *Srpk3-cKO* MZ B cells can differentiate into plasma cells in vitro.

Fig. S13. In vitro stimulated *Srpk3-cKO* MZ B cells can undergo class switch recombination.

Fig. S14. Purity of FACS sorted MZ B cells and relative variation between RNA-seq replicates across conditions.

Fig. S15. SRPK3 is necessary for regulating alternative splicing patterns in naïve and 72-hour stimulated MZ B cells.

Data file S1. Tpms values for immature and mature BM B cell populations between *WT* and *Srpk3*-deficient samples. Raw data.

Data file S2. Significant alternative splicing events between *WT* and *Srpk3*-deficient immature and mature BM B cells. Raw data.

Data file S3. Tpms of naïve, 8 hour and 72 hour stimulated *WT* and *Srpk3*-deficient marginal zone B cell samples. Raw data.

Data file S4. Significant alternative splicing events in naïve, 8-hour and 72-hour stimulated *WT* and *Srpk3*-deficient MZ B cells. Raw data.

Table S1. PCR Primer List

## Acknowledgments

We would like to thank J. Cambier for the gift of the *Cd79a-Cre^Tg/+^* mice. We thank L. Reinhardt and R. Kedl for the rmIL-5 and Seahorse reagents, respectively. We are grateful to R. Pelanda and C. Fleenor for helpful scientific discussions, M. Dell’Aringa for critically reading the paper, P. Strauch, J. Klarquist, T. Pfeifer, and J. Loomis for excellent technical assistance.

## Author contributions

Research conceptualization: TA, RMT and JRH. Performed research: TA, JMT, and EP. RNA-seq processing: JMT. Statistical analysis: TA, JMT, and JRK. Analyzed data: TA, JMT, EP, and JRK. Edited the paper: TA, JMT, EP, JRK, RMT, and JRH. Wrote the paper: TA, JMT, RMT, and JRH.

## Data and materials availability

The RNA-seq data have been deposited in the NCBI Gene Expression Omnibus (GEO; http://www.ncbi.nlm.nih.gov/geo/) repository under reference series GSE136622.

## References

1. A. Schaub, E. Glasmacher, Splicing in immune cells-mechanistic insights and emerging topics. Int Immunol 29, 173–181 (2017).

2. M. Yabas, H. Elliott, G. F. Hoyne, The Role of Alternative Splicing in the Control of Immune Homeostasis and Cellular Differentiation. Int J Mol Sci 17, (2015).

3. S. R. Bruce, B-cell and plasma-cell splicing differences: A potential role in regulated immunoglobulin RNA processing. Rna 9, 1264–1273 (2003).

4. A. M. Nelson, N. T. Carew, S. M. Smith, C. Milcarek, RNA Splicing in the Transition from B Cells to Antibody-Secreting Cells: The Influences of ELL2, Small Nuclear RNA, and Endoplasmic Reticulum Stress. J Immunol 201, 3073–3083 (2018).

5. W. P. Galej, Structural studies of the spliceosome: past, present and future perspectives. Biochem Soc Trans 46, 1407–1422 (2018).

6. S. D. Wagner, J. A. Berglund, Alternative pre-mRNA splicing. Methods Mol Biol 1126, 45–54 (2014).

7. J. F. Gui, W. S. Lane, X. D. Fu, A serine kinase regulates intracellular localization of splicing factors in the cell cycle. Nature 369, 678–682 (1994).

8. X. Y. Zhong, J. H. Ding, J. A. Adams, G. Ghosh, X. D. Fu, Regulation of SR protein phosphorylation and alternative splicing by modulating kinetic interactions of SRPK1 with molecular chaperones. Genes Dev 23, 482–495 (2009).

9. T. Giannakouros, E. Nikolakaki, I. Mylonis, E. Georgatsou, Serine-arginine protein kinases: a small protein kinase family with a large cellular presence. FEBS J 278, 570–586 (2011).

10. M. Varjosalo, S. Keskitalo, A. Van Drogen, H. Nurkkala, A. Vichalkovski, R. Aebersold, M. Gstaiger, The protein interaction landscape of the human CMGC kinase group. Cell Rep 3, 1306–1320 (2013).

11. J. F. Gui, H. Tronchere, S. D. Chandler, X. D. Fu, Purification and characterization of a kinase specific for the serine- and arginine-rich pre-mRNA splicing factors. Proc Natl Acad Sci U S A 91, 10824–10828 (1994).

12. H. Y. Wang, W. Lin, J. A. Dyck, J. M. Yeakley, Z. Songyang, L. C. Cantley, X. D. Fu, SRPK2: a differentially expressed SR protein-specific kinase involved in mediating the interaction and localization of pre-mRNA splicing factors in mammalian cells. J Cell Biol 140, 737–750 (1998).

13. J. Koizumi, Y. Okamoto, H. Onogi, A. Mayeda, A. R. Krainer, M. Hagiwara, The subcellular localization of SF2/ASF is regulated by direct interaction with SR protein kinases (SRPKs). J Biol Chem 274, 11125–11131 (1999).

14. Z. Zhou, J. Qiu, W. Liu, Y. Zhou, R. M. Plocinik, H. Li, Q. Hu, G. Ghosh, J. A. Adams, M. G. Rosenfeld, X. D. Fu, The Akt-SRPK-SR axis constitutes a major pathway in transducing EGF signaling to regulate alternative splicing in the nucleus. Mol Cell 47, 422–433 (2012).

15. D. Tsianou, E. Nikolakaki, A. Tzitzira, S. Bonanou, T. Giannakouros, E. Georgatsou, The enzymatic activity of SR protein kinases 1 and 1a is negatively affected by interaction with scaffold attachment factors B1 and 2. FEBS J 276, 5212–5227 (2009).

16. S. K. Petersen-Mahrt, C. Estmer, C. Ohrmalm, D. A. Matthews, W. C. Russell, G. Akusjarvi, The splicing factor-associated protein, p32, regulates RNA splicing by inhibiting ASF/SF2 RNA binding and phosphorylation. EMBO J 18, 1014–1024 (1999).

17. O. Nakagawa, M. Arnold, M. Nakagawa, H. Hamada, J. M. Shelton, H. Kusano, T. M. Harris, G. Childs, K. P. Campbell, J. A. Richardson, I. Nishino, E. N. Olson, Centronuclear myopathy in mice lacking a novel muscle-specific protein kinase transcriptionally regulated by MEF2. Genes Dev 19, 2066–2077 (2005).

18. P. R. Wilker, M. Kohyama, M. M. Sandau, J. C. Albring, O. Nakagawa, J. J. Schwarz, K. M. Murphy, Transcription factor Mef2c is required for B cell proliferation and survival after antigen receptor stimulation. Nat Immunol 9, 603–612 (2008).

19. I. D. R. Revilla, I. Bilic, B. Vilagos, H. Tagoh, A. Ebert, I. M. Tamir, L. Smeenk, J. Trupke, A. Sommer, M. Jaritz, M. Busslinger, The B-cell identity factor Pax5 regulates distinct transcriptional programmes in early and late B lymphopoiesis. EMBO J 31, 3130–3146 (2012).

20. I. Debnath, K. M. Roundy, P. D. Pioli, J. J. Weis, J. H. Weis, Bone marrow-induced Mef2c deficiency delays B-cell development and alters the expression of key B-cell regulatory proteins. Int Immunol 25, 99–115 (2013).

21. E. Hobeika, S. Thiemann, B. Storch, H. Jumaa, P. J. Nielsen, R. Pelanda, M. Reth, Testing gene function early in the B cell lineage in mb1-cre mice. Proc Natl Acad Sci U S A 103, 13789–13794 (2006).

22. R. Pelanda, E. Hobeika, T. Kurokawa, Y. Zhang, S. Kuppig, M. Reth, Cre recombinase-controlled expression of the mb-1 allele. Genesis 32, 154–157 (2002).

23. R. R. Hardy, P. W. Kincade, K. Dorshkind, The protean nature of cells in the B lymphocyte lineage. Immunity 26, 703–714 (2007).

24. J. Seita, D. Sahoo, D. J. Rossi, D. Bhattacharya, T. Serwold, M. A. Inlay, L. I. Ehrlich, J. W. Fathman, D. L. Dill, I. L. Weissman, Gene Expression Commons: an open platform for absolute gene expression profiling. PLoS One 7, e40321 (2012).

25. A. Cariappa, C. Chase, H. Liu, P. Russell, S. Pillai, Naive recirculating B cells mature simultaneously in the spleen and bone marrow. Blood 109, 2339–2345 (2007).

26. G. Huang, G. Ge, D. Wang, B. Gopalakrishnan, D. H. Butz, R. J. Colman, A. Nagy, D. S. Greenspan, alpha3(V) collagen is critical for glucose homeostasis in mice due to effects in pancreatic islets and peripheral tissues. J Clin Invest 121, 769–783 (2011).

27. M. N. Svensson, K. M. Andersson, C. Wasen, M. C. Erlandsson, M. Nurkkala-Karlsson, I. M. Jonsson, M. Brisslert, M. Bemark, M. I. Bokarewa, Murine germinal center B cells require functional Fms-like tyrosine kinase 3 signaling for IgG1 class-switch recombination. Proc Natl Acad Sci U S A 112, E6644–6653 (2015).

28. A. Y. Kreins, M. J. Ciancanelli, S. Okada, X. F. Kong, N. Ramirez-Alejo, S. S. Kilic, J. El Baghdadi, S. Nonoyama, S. A. Mahdaviani, F. Ailal, A. Bousfiha, D. Mansouri, E. Nievas, C. S. Ma, G. Rao, A. Bernasconi, H. Sun Kuehn, J. Niemela, J. Stoddard, P. Deveau, A. Cobat, S. El Azbaoui, A. Sabri, C. K. Lim, M. Sundin, D. T. Avery, R. Halwani, A. V. Grant, B. Boisson, D. Bogunovic, Y. Itan, M. Moncada-Velez, R. Martinez-Barricarte, M. Migaud, C. Deswarte, L. Alsina, D. Kotlarz, C. Klein, I. Muller-Fleckenstein, B. Fleckenstein, V. Cormier-Daire, S. Rose-John, C. Picard, L. Hammarstrom, A. Puel, S. Al-Muhsen, L. Abel, D. Chaussabel, S. D. Rosenzweig, Y. Minegishi, S. G. Tangye, J. Bustamante, J. L. Casanova, S. Boisson-Dupuis, Human TYK2 deficiency: Mycobacterial and viral infections without hyper-IgE syndrome. J Exp Med 212, 1641–1662 (2015).

29. R. Potla, T. Koeck, J. Wegrzyn, S. Cherukuri, K. Shimoda, D. P. Baker, J. Wolfman, S. M. Planchon, C. Esposito, B. Hoit, J. Dulak, A. Wolfman, D. Stuehr, A. C. Larner, Tyk2 tyrosine kinase expression is required for the maintenance of mitochondrial respiration in primary pro-B lymphocytes. Mol Cell Biol 26, 8562–8571 (2006).

30. C. L. Nguyen, R. Possemato, E. L. Bauerlein, A. Xie, R. Scully, W. C. Hahn, Nek4 regulates entry into replicative senescence and the response to DNA damage in human fibroblasts. Mol Cell Biol 32, 3963–3977 (2012).

31. Z. Zhou, X. D. Fu, Regulation of splicing by SR proteins and SR protein-specific kinases. Chromosoma 122, 191–207 (2013).

32. D. Dominguez, P. Freese, M. S. Alexis, A. Su, M. Hochman, T. Palden, C. Bazile, N. J. Lambert, E. L. Van Nostrand, G. A. Pratt, G. W. Yeo, B. R. Graveley, C. B. Burge, Sequence, Structure, and Context Preferences of Human RNA Binding Proteins. Mol Cell 70, 854–867 e859 (2018).

33. J. Xin, A. Mark, C. Afrasiabi, G. Tsueng, M. Juchler, N. Gopal, G. S. Stupp, T. E. Putman, B. J. Ainscough, O. L. Griffith, A. Torkamani, P. L. Whetzel, C. J. Mungall, S. D. Mooney, A. I. Su, C. Wu, High-performance web services for querying gene and variant annotation. Genome Biol 17, 91 (2016).

34. J. Wang, R. Al-Ouran, Y. Hu, S. Y. Kim, Y. W. Wan, M. F. Wangler, S. Yamamoto, H. T. Chao, A. Comjean, S. E. Mohr, Udn, N. Perrimon, Z. Liu, H. J. Bellen, MARRVEL: Integration of Human and Model Organism Genetic Resources to Facilitate Functional Annotation of the Human Genome. Am J Hum Genet 100, 843–853 (2017).

35. F. Martin, J. F. Kearney, Marginal-zone B cells. Nat Rev Immunol 2, 323–335 (2002).

36. L. M. Pecanha, C. M. Snapper, F. D. Finkelman, J. J. Mond, Dextran-conjugated anti-Ig antibodies as a model for T cell-independent type 2 antigen-mediated stimulation of Ig secretion in vitro. I. Lymphokine dependence. J Immunol 146, 833–839 (1991).

37. C. M. Snapper, T. M. McIntyre, R. Mandler, L. M. Pecanha, F. D. Finkelman, A. Lees, J. J. Mond, Induction of IgG3 secretion by interferon gamma: a model for T cell-independent class switching in response to T cell-independent type 2 antigens. J Exp Med 175, 1367–1371 (1992).

38. M. J. McCarron, P. W. Park, D. R. Fooksman, CD138 mediates selection of mature plasma cells by regulating their survival. Blood 129, 2749–2759 (2017).

39. J. Tellier, S. L. Nutt, Standing out from the crowd: How to identify plasma cells. Eur J Immunol 47, 1276–1279 (2017).

40. M. Scalise, L. Pochini, L. Console, M. A. Losso, C. Indiveri, The Human SLC1A5 (ASCT2) Amino Acid Transporter: From Function to Structure and Role in Cell Biology. Front Cell Dev Biol 6, 96 (2018).

41. L. D. Osellame, A. P. Singh, D. A. Stroud, C. S. Palmer, D. Stojanovski, R. Ramachandran, M. T. Ryan, Cooperative and independent roles of the Drp1 adaptors Mff, MiD49 and MiD51 in mitochondrial fission. J Cell Sci 129, 2170–2181 (2016).

42. J. A. Ronchi, T. R. Figueira, F. G. Ravagnani, H. C. Oliveira, A. E. Vercesi, R. F. Castilho, A spontaneous mutation in the nicotinamide nucleotide transhydrogenase gene of C57BL/6J mice results in mitochondrial redox abnormalities. Free Radic Biol Med 63, 446–456 (2013).

43. L. Liu, C. Luo, Y. Luo, L. Chen, Y. Liu, Y. Wang, J. Han, Y. Zhang, N. Wei, Z. Xie, W. Wu, G. Wu, Y. Feng, MRPL33 and its splicing regulator hnRNPK are required for mitochondria function and implicated in tumor progression. Oncogene 37, 86–94 (2018).

44. B. K. Chacko, P. A. Kramer, S. Ravi, G. A. Benavides, T. Mitchell, B. P. Dranka, D. Ferrick, A. K. Singal, S. W. Ballinger, S. M. Bailey, R. W. Hardy, J. Zhang, D. Zhi, V. M. Darley-Usmar, The Bioenergetic Health Index: a new concept in mitochondrial translational research. Clin Sci (Lond) 127, 367–373 (2014).

45. J. Han, J. H. Ding, C. W. Byeon, J. H. Kim, K. J. Hertel, S. Jeong, X. D. Fu, SR proteins induce alternative exon skipping through their activities on the flanking constitutive exons. Mol Cell Biol 31, 793–802 (2011).

46. T. Bradley, M. E. Cook, M. Blanchette, SR proteins control a complex network of RNA-processing events. RNA 21, 75–92 (2015).

47. S. T. Ju, M. E. Dorf, Preferential induction of specific lambda-isotypic antibodies in mice. J Immunol 133, 1404–1409 (1984).

48. F. Melchers, Checkpoints that control B cell development. J Clin Invest 125, 2203–2210 (2015).

49. G. E. Woloschak, C. J. Krco, Regulation of kappa/lambda immunoglobulin light chain expression in normal murine lymphocytes. Mol Immunol 24, 751–757 (1987).

50. J. J. Bednarski, B. P. Sleckman, Lymphocyte development: integration of DNA damage response signaling. Adv Immunol 116, 175–204 (2012).

51. J. J. Bednarski, R. Pandey, E. Schulte, L. S. White, B. R. Chen, G. J. Sandoval, M. Kohyama, M. Haldar, A. Nickless, A. Trott, G. Cheng, K. M. Murphy, C. H. Bassing, J. E. Payton, B. P. Sleckman, RAG-mediated DNA double-strand breaks activate a cell type-specific checkpoint to inhibit pre-B cell receptor signals. J Exp Med 213, 209–223 (2016).

52. J. Hu, S. Tepsuporn, R. M. Meyers, M. Gostissa, F. W. Alt, Developmental propagation of V(D)J recombination-associated DNA breaks and translocations in mature B cells via dicentric chromosomes. Proc Natl Acad Sci U S A 111, 10269–10274 (2014).

53. E. Jacque, E. Schweighoffer, V. L. Tybulewicz, S. C. Ley, BAFF activation of the ERK5 MAP kinase pathway regulates B cell survival. J Exp Med 212, 883–892 (2015).

54. M. Ota, B. H. Duong, A. Torkamani, C. M. Doyle, A. L. Gavin, T. Ota, D. Nemazee, Regulation of the B cell receptor repertoire and self-reactivity by BAFF. J Immunol 185, 4128–4136 (2010).

55. T. N. Iwata, J. A. Ramirez-Komo, H. Park, B. M. Iritani, Control of B lymphocyte development and functions by the mTOR signaling pathways. Cytokine Growth Factor Rev 35, 47–62 (2017).

56. M. J. Gros, P. Naquet, R. R. Guinamard, Cell intrinsic TGF-beta 1 regulation of B cells. J Immunol 180, 8153–8158 (2008).

57. K. E. Gunn, J. W. Brewer, Evidence that marginal zone B cells possess an enhanced secretory apparatus and exhibit superior secretory activity. J Immunol 177, 3791–3798 (2006).

58. F. Martin, A. M. Oliver, J. F. Kearney, Marginal zone and B1 B cells unite in the early response against T-independent blood-borne particulate antigens. Immunity 14, 617–629 (2001).

59. N. S. Greenspan, L. J. Cooper, Cooperative binding by mouse IgG3 antibodies: implications for functional affinity, effector function, and isotype restriction. Springer Semin Immunopathol 15, 275–291 (1993).

60. M. C. Hsu, K. M. Toellner, C. G. Vinuesa, I. C. Maclennan, B cell clones that sustain long-term plasmablast growth in T-independent extrafollicular antibody responses. Proc Natl Acad Sci U S A 103, 5905–5910 (2006).

61. N. Degauque, C. Brosseau, S. Brouard, Regulation of the Immune Response by the Inflammatory Metabolic Microenvironment in the Context of Allotransplantation. Front Immunol 9, 1465 (2018).

62. M. Muschen, Metabolic gatekeepers to safeguard against autoimmunity and oncogenic B cell transformation. Nat Rev Immunol 19, 337–348 (2019).

63. C. A. Doughty, B. F. Bleiman, D. J. Wagner, F. J. Dufort, J. M. Mataraza, M. F. Roberts, T. C. Chiles, Antigen receptor-mediated changes in glucose metabolism in B lymphocytes: role of phosphatidylinositol 3-kinase signaling in the glycolytic control of growth. Blood 107, 4458–4465 (2006).

64. L. R. Waters, F. M. Ahsan, D. M. Wolf, O. Shirihai, M. A. Teitell, Initial B Cell Activation Induces Metabolic Reprogramming and Mitochondrial Remodeling. iScience 5, 99–109 (2018).

65. E. Krzywinska, C. Stockmann, Hypoxia, Metabolism and Immune Cell Function. Biomedicines 6, (2018).

66. P. A. Gameiro, L. A. Laviolette, J. K. Kelleher, O. Iliopoulos, G. Stephanopoulos, Cofactor balance by nicotinamide nucleotide transhydrogenase (NNT) coordinates reductive carboxylation and glucose catabolism in the tricarboxylic acid (TCA) cycle. J Biol Chem 288, 12967–12977 (2013).

67. M. D. Brand, D. G. Nicholls, Assessing mitochondrial dysfunction in cells. Biochem J 435, 297–312 (2011).

68. J. M. Son, E. H. Sarsour, A. Kakkerla Balaraju, J. Fussell, A. L. Kalen, B. A. Wagner, G. R. Buettner, P. C. Goswami, Mitofusin 1 and optic atrophy 1 shift metabolism to mitochondrial respiration during aging. Aging Cell 16, 1136–1145 (2017).

69. D. P. Corkery, A. C. Holly, S. Lahsaee, G. Dellaire, Connecting the speckles: Splicing kinases and their role in tumorigenesis and treatment response. Nucleus 6, 279–288 (2015).

70. K. Lukin, S. Fields, D. Lopez, M. Cherrier, K. Ternyak, J. Ramirez, A. J. Feeney, J. Hagman, Compound haploinsufficiencies of Ebf1 and Runx1 genes impede B cell lineage progression. Proc Natl Acad Sci U S A 107, 7869–7874 (2010).

71. C. L. Swanson, T. J. Wilson, P. Strauch, M. Colonna, R. Pelanda, R. M. Torres, Type I IFN enhances follicular B cell contribution to the T cell-independent antibody response. J Exp Med 207, 1485–1500 (2010).

72. M. Martin. Cutadapt Removes Adapter Sequences From High-Throughput Sequencing Reads. EMBnet.journal 17, 10–12 (2011).

73. R. Patro, G. Duggal, M. I. Love, R. A. Irizarry, C. Kingsford, Salmon provides fast and bias-aware quantification of transcript expression. Nat Methods 14, 417–419 (2017).

74. C. Soneson, M. I. Love, M. D. Robinson, Differential analyses for RNA-seq: transcript-level estimates improve gene-level inferences. F1000Res 4, 1521 (2015).

75. M. I. Love, W. Huber, S. Anders, Moderated estimation of fold change and dispersion for RNA-seq data with DESeq2. Genome Biol 15, 550 (2014).

76. A. Dobin, C. A. Davis, F. Schlesinger, J. Drenkow, C. Zaleski, S. Jha, P. Batut, M. Chaisson, T. R. Gingeras, STAR: ultrafast universal RNA-seq aligner. Bioinformatics 29, 15–21 (2013).

77. S. Shen, J. W. Park, Z. X. Lu, L. Lin, M. D. Henry, Y. N. Wu, Q. Zhou, Y. Xing, rMATS: robust and flexible detection of differential alternative splicing from replicate RNA-Seq data. Proc Natl Acad Sci U S A 111, E5593–5601 (2014).

