## Supplementary Materials for "SRPK3 regulates alternative pre-mRNA splicing required for B lymphocyte development and humoral responsiveness"

Materials and Methods

*Antibodies*

The following antibodies were used for flow cytometry: anti-Ly6c Phycoerythrin (PE) (HK1.4, eBiosciences), anti-NK1.1 PE (PK136, eBiosciences), anti-CD317 PE (eBio927, eBiosciences), anti-CD11b (various fluorochromes, M1/70, Invitrogen, Biolegend), anti-CD11c PE (N418, Invitrogen), anti-Ter119 PE (BD Biosciences), anti-Gr1 PE (RB6-8C5, eBiosciences), anti-CD3ε (various fluorochromes, 145-2C11, BD Biosciences), $NP_{28}$ PE (LGC Biosearch Technologies), anti-CD23 (various fluorochromes, B3B4, eBiosciences, BD Biosciences), anti-Ig lambda (various fluorochromes, RML-42, Biolegend, Southern Biotech), anti-Ig kappa (various fluorochromes, 187.1, BD Biosciences, Pharmingen), anti-TACI PE (8F10, Biolegend), anti-B220 (various fluorochromes, RA3-6B2, Biolegend, eBiosciences) anti-CD19 (various fluorochomes, 6D5, Biolegend, 1D3, eBiosciences, BD Biosciences), anti-CD25 Allophycocyanin (APC)-Cy7 (PC61, Biolegend), anti-CD117 APC (2B8, eBiosciences), anti-CD93 APC (AA4.1, eBiosciences), anti-IgD (various fluorochromes, 11-26C.2a, Biolegend), anti-CD21/35 (various fluorochromes, 7E9, Biolegend, 7G6, BD Biosciences), anti-CD125 APC (Miltenyi Biotech), Streptavidin APC (BD Pharmingen), anti-IgM (various fluorochromes, RMM-1, Biolegend), anti-CD119 Biotin (2E2, Invitrogen), anti-CD43 Fluorescein isothiocyanate (FITC) (S7, BD Biosciences), anti-CD24 (various fluorochromes, 30-F1 or M1-69, eBiosciences), anti-BP1 Peridinin Chlorophyll Protein Complex (PerCP) e710 (6C3, Invitrogen), and anti-CD138 Brilliant Violet (BV) 421 (281-2, Biolegend). The bone marrow $Lin^-$ cocktail consisted of antibodies (listed above) CD3ε, CD11b, CD11c, Gr-1. NK1.1, Ly6c, CD317, and Ter119. The spleen $Lin^-$ consisted of CD3ε and CD11b. A Live/Dead Fixable Violet

Dead Cell stain was used for viability analysis according to manufacturer's instructions (L34955, Life Technologies).

*Cell preparation and flow cytometry*

For FACS analysis, bone marrow or spleens of adult mice were harvested in Growth media [RPMI (Life Technologies) supplemented with 1X Glutamax (Life Technologies), 50µg/mL gentamicin (Life Technologies), 50µM β-mercaptoethanol (Sigma-aldrich) and 10% FBS (HyClone)]. Red blood cells (RBCs) were lysed in 2 mL of Hemolytic buffer (0.155M $NH_4Cl$, 0.01M $NaHCO_3$, 0.1mM $Na_2$ EDTA) for 3 min on ice and quenched with 20 mL Growth media. Cells were washed with 10mL FACS buffer (1X PBS (pH 7.6), 5% FBS, 4mM EDTA) and stained in FACS buffer containing antibodies and $F_c$ block (anti-CD16/CD32; eBioscience) for at least 30 min on ice. Cells were subsequently washed 3X with FACS buffer and analyzed. Characterization of phenotypic fractions included pre-pro B cells (Fraction A; Lin$^-$B220$^+$CD43$^+$BP1$^-$CD19$^-$), pro B cells (Fraction B/C; Lin$^-$B220$^+$CD43$^+$BP1$^-$CD19$^+$), early pre B cells (Fraction C'; Lin$^-$B220$^+$CD43$^+$BP1$^+$CD19$^+$), late pre B cells (Fraction D; Lin$^-$B220$^+$CD43$^-$CD24$^+$IgM$^-$BP1$^+$), immature B cells (Fraction E; Lin$^-$B220$^+$CD43$^-$CD93$^+$IgM$^+$IgD$^-$) and recirculating mature B cells (Fraction F; Lin$^-$B220$^+$CD43$^-$CD93$^-$IgM$^+$IgD$^+$). For cell sort preparation, bone marrow or spleens of adult mice were harvested in Growth media. RBCs were lysed in 2 mL of Hemolytic buffer for 3 min on ice and quenched with 20 mL Growth media. Cells were washed with 10mL MoFlow buffer (1X PBS (pH 7.6), 2% FBS) and stained in MoFlow buffer containing antibodies and $F_c$ block for at least 30 min on ice. Cells were subsequently washed 3X with MoFlow buffer and analyzed. All cells were gated on live, singlet populations. LSR II (BD Biosciences), LSR Fortessa (BD Biosciences), or Synergy (iCyt) cytometers were used for flow cytometry analysis or cell sorting. Data were analyzed using FlowJo v9.2 or v10 software (TreeStar).

*Intracellular flow cytometry*

Intracellular flow cytometry was performed using the BrdU flow kit (BD Biosciences). In brief, bone marrow cells were harvested and stained for cell surface antigens. Cells were then fixed and permeabilized with BD Cytofix/Cytoperm buffer for 15 min at room temperature (RT). Cells were washed with 1X BD Perm/Wash buffer. Cells were then incubated with BD Cytoperm Permeabilization buffer for 10 min on ice and washed. Cells were re-fixed with Cytofix/Cytoperm buffer for 5 min at RT and washed. Cells were then stained with intracellular antigens and incubated 15 min at RT, washed and analyzed.

*Immature and mature bone marrow B cell library preparation and sequencing*

Immature B cells ($B220^{+}Lin^{-}CD19^{+}CD25^{-}CD43^{-}CD93^{+}$) and mature B cells ($B220^{+}Lin^{-}CD19^{+}CD25^{-}CD43^{-}CD93^{-}$) were isolated from bone marrow of *Srpk3*$^{fl}$*Cd79a-Cre*$^{Tg/+}$ *(Srpk3-cKO)* or *WT* littermate controls using the Synergy Sorter (iCyt). Purity of sorted cells from *Srpk3-cKO* and *WT* mice was determined by back-sort using Synergy Sorter (iCyt) (Fig. S5A). RNA from these cells (~700,000-3,000,000 total cells per biological replicate) was isolated using the RNeasy Micro kit (Qiagen), with on-column treatment with RNase-free DNaseI, according to manufacturer’s instructions. RNA analysis, cDNA synthesis and library preparation were performed by the Genomics and Microarray Core facility at the University of Colorado Anschutz Medical Campus (UCD-AMC). The Agilent Tape Station 4200 was used to assess RNA quality. Poly A selected sequencing libraries were constructed from 100ng of Total RNA using the Nugen Universal Plus mRNA –SEQ kit (Tecan). Libraries were quantitated using Qubit and assessed on the Agilent Tape Station 4200 using a DNA 1000 screen tape. Sequencing was done with 2x150bp chemistry on the Illumina NovaSEQ6000 Instrument using an S4 flow cell.

*Enzyme-linked immunosorbent assay*

For regular ELISA assays, 96-well plates were coated with IgH+L for 1 hour at 37°C. All steps were performed in a humid chamber. Wells were washed 3X with Wash buffer (1X PBS, 0.1% Tween). Wells were blocked with Blocking buffer (1X PBS, 1% BSA) overnight (O/N) at 37°C and washed with Wash buffer. Wells were incubated with Ig standards (IgM, IgG1, IgG3, IgA, IgG2c) and mouse serum for 1 hour at 37°C and washed with Wash buffer. Wells were then incubated with polyclonal goat anti-mouse AP-conjugated antibodies for 1 hour at 37°C and then washed with Wash buffer. Wells were allowed to develop with phosphatase substrate (Sigma) in Develop buffer (0.1M Diethanolamine, 8mM $MgCl_2$, 0.02% $NaN_3$, pH 9.8) for at least 30 min at 37°C. Plates were read at OD405 on the Versamax Microplate reader (Molecular Devices). For NP-specific ELISA assays, all steps remained the same as regular ELISA unless otherwise noted and previously described (*71*). 96-well plates were coated with $NIP_{(22)}$-BSA for 1 hour at 37°C. Wells were incubated with NP-Ig standards (IgM (B18μ), IgG1 (N1G9), IgG3 (S24/63/63) and mouse serum for 1 hour at 37°C and washed with Wash buffer.

*Immunization of mice*

Adult mice were immunized with intraperitoneal injections of either 200µL 25µg/mL $NP_{40}$-AECM-FICOLL (LGC Biosearch Technologies), 200µL 0.5mg/mL NP-CGG (ratio 30-39; LGC Biosearch Technologies), 200µL 100µg/mL $NP\text{-}LPS_{(0.3)}$ (LGC Biosearch Techonolgies) or 200µL 1X PBS (control). Mice were bled (100µL) from the submandibular vein prior to immunization and every 7 days post-injection (p.i.) until day 56 p.i. Serum from blood was extracted using Micro tube 1.1mL Z-gel (Sarstedt). In addition, mice were euthanized on day 4, 6, or 14 p.i. for flow cytometric analysis of NP-specific populations or for qRT-PCR analysis.

*Class switch recombination qRT-PCR*

For immunized mice, B cells ($CD19^+Lin^-$) were isolated from spleens of NP-Ficoll immunized *Srpk3-cKO* or *WT* littermate controls using the Synergy Sorter (iCyt). RNA from these cells (500,000 total cells per biological replicate) was isolated and cDNA was synthesized. qRT-PCR was ran using the AB7300 (Applied Biosystems). Technical triplicates were done for all biological replicates. Primers used can be found in Table S1. For vitro culture, RNA from cultured cells (100,000 total cells per biological replicate) was isolated and cDNA was synthesized. qRT-PCR was ran using the QuantStudio5 (Thermo Fisher). Technical triplicates were done for all biological replicates.

*Marginal zone B cell in vitro stimulation*

Marginal zone B cells ($Lin^-B220^+CD21^{hi}CD24^{hi}CD23^-CD19^+$) were isolated from spleens of *Srpk3-cKO* or *WT* littermate controls using the Synergy Sorter (iCyt). Cells were plated at $1.0x10^5$ cells/mL in Growth media. Cells were stimulated with 3ng/mL α-δ-dex (FinaBio), 10ng/mL rmIL-5 (R&D Systems) and 10U/mL rmIFN-γ (Biolegend). Cells remained in culture for 8 hours, 24 hours, 72 hours or 6 days. For cells in culture for 72 hours or 6 days, IFN-γ was added after 24 hours pre-stimulation.

*Marginal Zone B cell library preparation and RNA-sequencing*

Marginal zone B cells ($Lin^-B220^+CD21^{hi}CD24^{hi}CD23^-CD19^+$) were isolated from spleens of *Srpk3-cKO* or *WT* littermate controls using the Synergy Sorter (iCyt) and stimulated in vitro. Purity of sorted cells from *Srpk3-cKO* and *WT* mice was determined by back-sorting using Synergy Sorter (iCyt) (Fig. S13A). RNA from these cells (~50,000-150,000 total cells per biological replicate) was isolated using the RNeasy Micro kit (Qiagen), with on-column treatment with RNase-free DNaseI, according to manufacturer's instructions. RNA analysis, cDNA synthesis and library preparation were performed by the Genomics Core in the Center for Genes, Environment and Health at National Jewish Health. RNA concentration was assessed with the Qubit High Sensitivity RNA assay (Life Technologies) and quality was assessed with the Eukaryotic RNA Pico kit (Agilent) on the Bio Analyzer 2100. Isolated total RNA was processed for next-generation sequencing (NGS) library construction as developed in the NJH Genomics Facility for analysis with a HiSeq 2500 (San Diego, CA, USA). A Clontech SMART-Seq® v4 Ultra® Low Input RNA kit for sequencing (Mountain View, CA, USA) and Nextera XT (San Diego, CA, USA) kit were used. Briefly, library construction started from isolation of total RNA species, followed by SMARTer 1$^{st}$ strand cDNA synthesis, full length dscDNA amplification by LD-PCR, followed by purification and validation. After that, samples were taken to the Nextera XT protocol where the sample is simultaneously fragmented and tagged with adapters followed by a limited cycle PCR that adds indexes. Library concentration was assessed using Qubit High Sensitivity DNA assay (Agilent) and quality was assessed using the

High Sensitivity DNA kit (Agilent) on the Bioanalyzer 2100. Once validated, the barcoded-pooled libraries were sequenced using 2x125bp chemistry on the HiSeq 2500 as routinely performed by the NJH Genomics Facility.

*RNA-sequencing analysis and validation*

For sequencing analysis, reads were stripped of adapters from their 3' ends using cutadapt 1.9.1 (*72*). For gene expression analyses, transcript abundances were calculated against the mouse transcriptome (Gencode M17) using salmon 0.11 (*73*), including the flags —seqBias and —gcBias.  These quantifications were then collapsed to gene-level quantifications using tximport (*74*), and differentially expressed genes were identified using DESeq2 (*75*). The Wald test was used for statistical analyses for gene expression evaluation. That p value was then adjusted using the Benjamini-Hochsberg method. For splicing analyses, reads were aligned to the mouse genome (Gencode M17) using STAR 2.4.0 (*76*).  Alternative exon inclusion rates were calculated with rMATS (*77*).

The canonical pathway enrichment analyses were generated using Ingenuity Pathway Analysis (QIAGEN Inc., https://www.qiagenbio- informatics.com/products/ingenuity-pathway-analysis). A list of genes with a significant FDR ≤ 0.05 of alternative splicing event were uploaded to IPA. A Core Test was run with default settings except select all on confidence was chosen.

For RNA-seq validation, immature B cells ($B220^+Lin^-CD19^+CD25^-CD43^-CD93^+$) and mature B cells ($B220^+Lin^-CD19^+CD25^-CD43^-CD93^-$) were isolated from bone marrow of *Srpk3-cKO* or *WT* littermate controls using the Synergy Sorter (iCyt). RNA from these cells (~700,000-3,000,000 total cells per biological replicate) was isolated and cDNA synthesized. Marginal zone

B cells ($Lin^-B220^+CD21^{hi}CD24^{hi}CD23^-CD19^+$) were isolated from spleens of *Srpk3-cKO* or *WT* littermate controls using the Synergy Sorter (iCyt) and stimulated in vitro. RNA from these cells (~50,000-150,000 total cells per biological replicate) was isolated and cDNA synthesized. qRT-PCR was ran using the QuantStudio5 (Thermo Fisher). Primers used for validation can be found in Table S1. Technical triplicates were done for all biological replicates.

*Mitochondrial Mass*

Total splenocytes were isolated from *Srpk3-cKO* or *WT* littermate controls and plated in vitro in Growth media without stimulations for 2 hours or with 3ng/mL α-δ-dex (FinaBio), 10ng/mL rmIL-5 (R&D Systems) and 10U/mL rmIFN-γ (Biolegend) for 24 hours. Mitotracker Green FM (Thermo Fisher) was added directly to culture at a final concentration of 50nM for 30 min or DMSO was used as a control. Cells were then washed with FACS buffer, stained for cell surface markers and marginal zone B cells were then analyzed by flow cytometry.

*Structured Illumination Microscopy*

Marginal zone B cells ($Lin^-B220^+CD21^{hi}CD24^{hi}CD23^-CD19^+$) were isolated from spleens of *Srpk3-cKO* or *WT* littermate controls using the Synergy Sorter (iCyt). Cells were seeded onto glass bottom dishes (MaTek) pre-coated with 0.1mg/mL poly-D-lysine (Sigma) and incubated in Growth media in the presence of 10ng/mL rmBAFF (R&D Systems) only or BAFF with 3ng/mL α-δ-dex (FinaBio), 10ng/mL rmIL-5 (R&D Systems) and 10U/mL rmIFN-γ (Biolegend) for 24-hours. Cells were stained with Mitotracker Red (Thermo Fisher) at a final concentration of 100 nM for 15 minutes, washed, then imaged using a Nikon nSIM. Images shown are individual z-slices reconstructed using Nikon Elements with the 'Slice Reconstruction' tool.

*Metabolic flux assay*

Splenic B cells (CD43-) were isolated using MACS LS columns (Miltenyi Biotec) with anti-CD43 microbeads (Miltenyi Biotec) according to manufactures instructions. Cells were then plated with Growth media and left unstimulated or stimulated in vitro for 3 hours prior to analysis. Oxygen consumption rates (OCR) were determined using Seahorse XFe96 Analyzers (Agilent). Cells were seeded on 96-well plates (Agilent) pre-treated with poly-D-lysine (Sigma) at $4.0 \times 10^5$ cells per well in Seahorse XF RPMI media containing 2mM L-glutamine and 10mM dextrose. Plates were incubated for 1 hour at 37°C without $CO_2$ prior to OCR measurements. OCR was measured before and after injections of: 1μM oligomycin, 1μM carbonyl cyanide 4-(trifluoromethoxy) phenylhydrazone (FCCP), 4μM antimycin A, and 1μM rotenone. All concentrations listed are final concentrations of components. Basal respiration was determined by subtracting initial OCR values with OCR values after antimycin/rotenone administration. ATP-linked respiration was determined by subtracting basal OCR values with OCR values after oligomycin administration. Maximal respiration was determined by subtracting OCR values after FCCP administration with OCR values after antimycin/rotenone administration. Lastly, non-mitochondrial respiration were OCR values after antimycin/rotenone administration.

*Statistical analysis*

All statistical analyses were conducted using GraphPad Prism version 7.0 or 8.0. Each assay was repeated at least three times in independent experiments, unless noted otherwise. Mouse experiments were done using at least three mice per genotype per condition. Figure legends detail number of experimental replicates and *n* values. All data shown are means ± SEM, unless otherwise specified. All statistics shown were generated using unpaired t-tests, 1-way ANOVA or 2-way ANOVA unless otherwise specified. *p* values less than 0.05 were considered significant. On graphs, statistics are indicated with asterisks: * $p < 0.05$, ** $p < 0.005$. For RNA-seq, $\Delta\psi \geq 0.1$, FDR $< 0.05$. No overlap in $\psi$ between replicates across conditions.

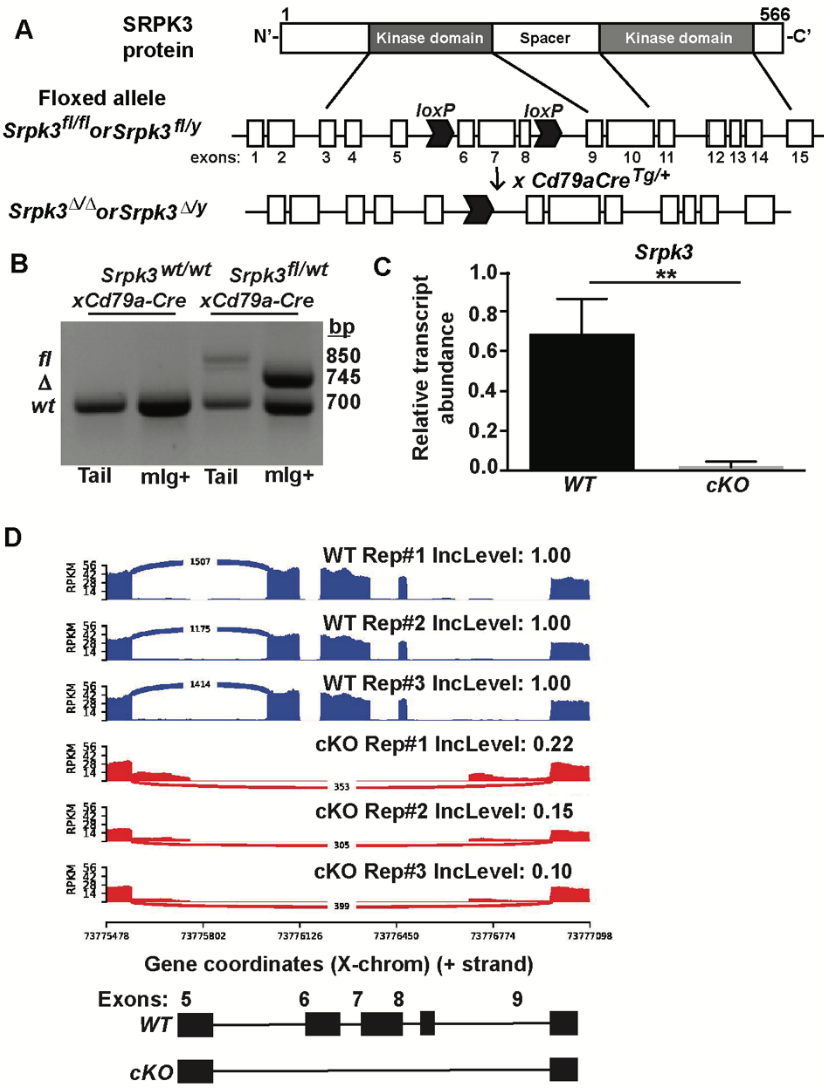


**Fig. S1. Efficient deletion of floxed *Srpk3* alleles in bone marrow and splenic B cells.** (**A**) Conditional inactivation of *Srpk3* in B cells. Diagram of *loxP* sites that flank *Srpk3* exons 6-8. *Cd79a*-Cre recombinase mediated deletion of floxed sites result in *Srpk3*$^{Δ/Δ}$ or *Srpk3*$^{Δ/y}$ allele with loss of amino acids 183-281 (X1 isoform). Joining of exons 5 and 9 results in a translational frameshift. (**B**) Deletion of *Srpk3* floxed allele in mIg+ bone marrow B cells sorted from *WT* or *Srpk3*$^{fl/WT}$*Cd79a-Cre*$^{Tg/+}$ (*Srpk3-hmz)* mice and tail DNA from the same mice. Relative quantification of transcripts with deleted exons were also confirmed by quantitative RT-PCR (qRT-PCR). Bone marrow mIg$^+$ B cells (Lin$^-$CD19$^+$B220$^+$CD43$^-$IgM$^+$) were sort purified using the Synergy Sorter (iCyt). RNA from these cells (~500,000 total cells per biological replicate) was isolated using the RNeasy Micro kit (Qiagen), with on-column treatment with RNase-free DNaseI, according to manufacturer's instructions. cDNA was synthesized according to SuperScript™ II Reverse Transcriptase (Thermo Fisher) protocol. qRT-PCR was ran using AB7300 (Applied Biosystems). Technical triplicates were performed for all biological replicates. (**C**) Relative expression of *Srpk3* in mIg+ bone marrow B cells sorted from *WT* or *Srpk3$^{fl/y}$Cd79a-Cre$^{Tg/+}$* (*Srpk3-cKO*) mice. Anti-sense primer spans portion of exon 6. (**D**) Sashimi plot of exon junction reads of *Srpk3* in marginal zone B cells purified from *WT* (n=3) or *Srpk3$^{fl/y}$Cd79a-Cre$^{Tg/+}$* (*Srpk3-cKO*) (n=3) male mice. Plot shows no reads spanning exons 6-8 in *cKO* cells. Asterisks indicate statistical significance compared to *WT* littermate controls as unpaired, two-tailed Student's t-test. **P<0.005. Graphs represent arithmetic mean ± SEM. All data are representative of at least three independent experiments.

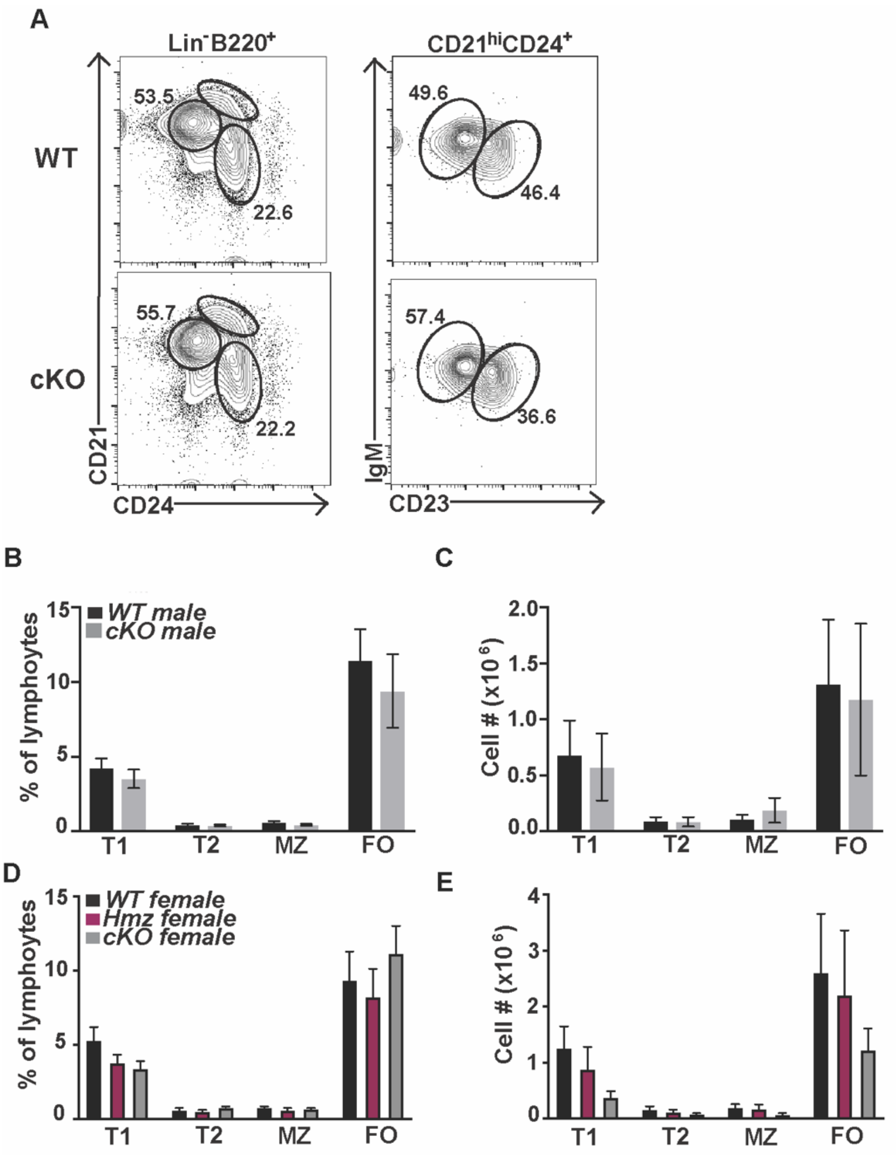


**Fig. S2. Normal splenic B cell development in mice that lack SRPK3.** (**A**) Phenotype, (**B** and **D**) frequencies and (**C** and **E**) cell numbers of various splenic B cell subsets between male *WT* (n=7-10) or *Srpk3-cKO* (n=7-9) mice (**B** and **C**) and female *WT* (n=5), *Srpk3-hemizygous* (*Hmz*, n=7), *Srpk3-cKO* (n=3) mice (**D** and **E**). In panel A numbers represent frequency of cells within phenotypic fractions. Splenic phenotypic subsets used were: Transitional 1 (T1; $Lin^-B220^+CD24^{hi}CD21^{lo}$), Transitional 2 (T2; $Lin^-B220^+CD24^{hi}CD21^{int}CD23^{hi}IgM^+$), Marginal zone (MZ; $Lin^-B220^+CD24^{hi}CD21^{hi}CD23^-IgM^+$) and Follicular (FO; $Lin^-B220^+CD24^{int}CD21^{int}$). 2-

way ANOVA was performed to determine statistical significance. Graphs represent arithmetic mean ± SEM. All data are representative of at least three independent experiments.

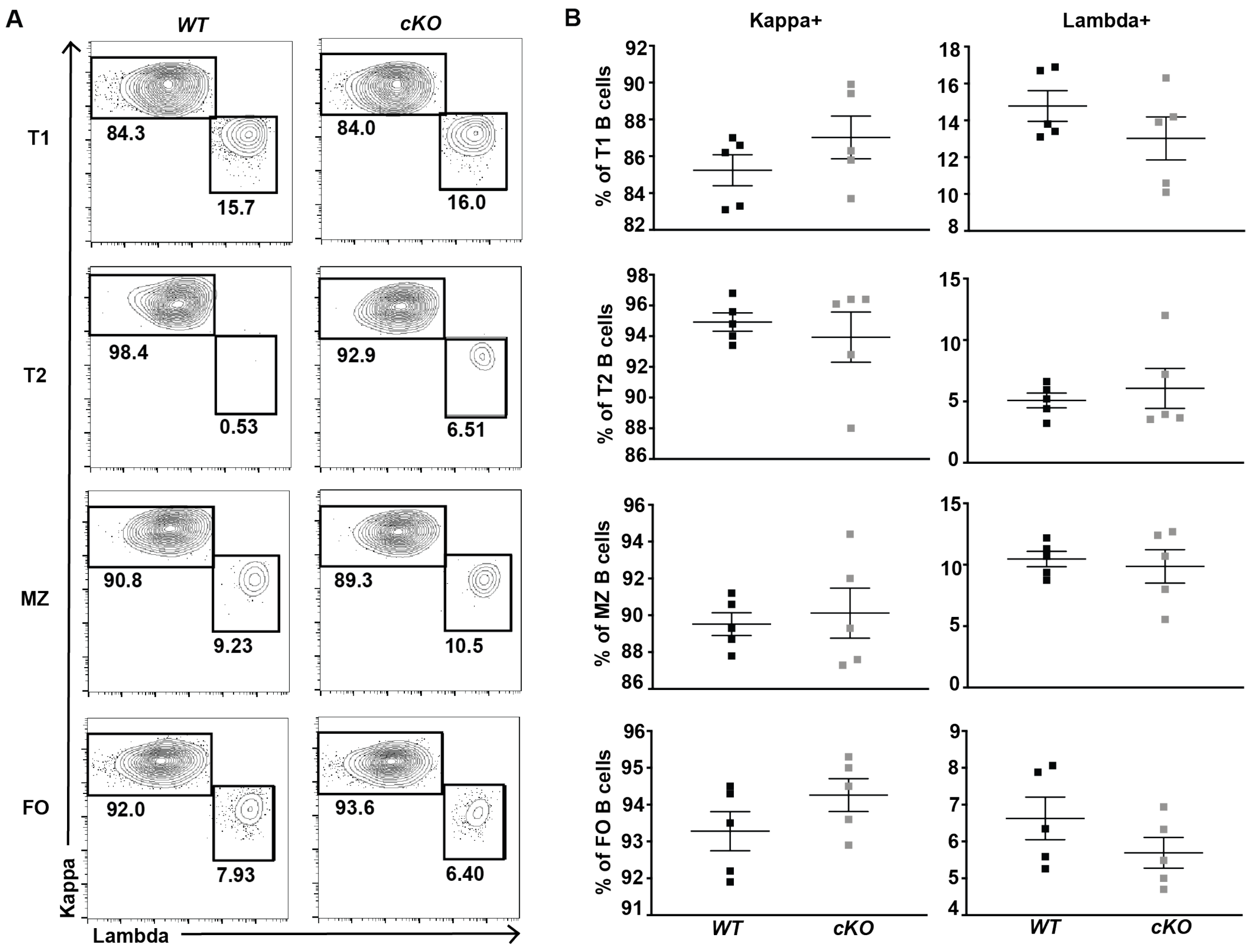


**Fig. S3. Kappa and lambda light chain expression on splenic B cell subsets in *Srpk3-cKO* mice.** (**A**) Flow cytometric analyses of kappa and lambda expression and (**B**) frequencies of kappa and lambda light chain expression on various splenic B cell subsets in *WT* (n=5) and *Srpk3-cKO* (n=5) male mice. T1 (Transitional 1), T2 (Transitional 2), MZ (Marginal Zone), FO (Follicular). Numbers in panel A refer to frequency of cells within phenotypic fraction. Two-tailed, Student's t-test was performed to determine statistical significance. Graphs represent arithmetic mean ± SEM. All data are representative of at least three independent experiments.

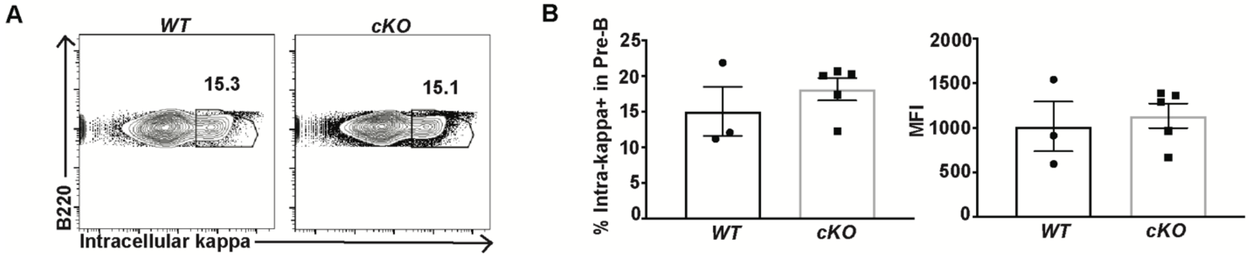


**Fig. S4. Intracellular kappa expression in bone marrow pre-B cells.** (**A**) Flow cytometric analyses and (**B**) frequencies and MFI of intracellular kappa expression in pre-B cells ($B220^{+}CD43^{-}IgM^{-}$) in *WT* (n=3) and *Srpk3-cKO* (n=5) male mice. Mann-Whitney Rank test was performed to determine statistical significance. Graphs represent arithmetic mean ± SEM. All data are representative of at least three independent experiments.

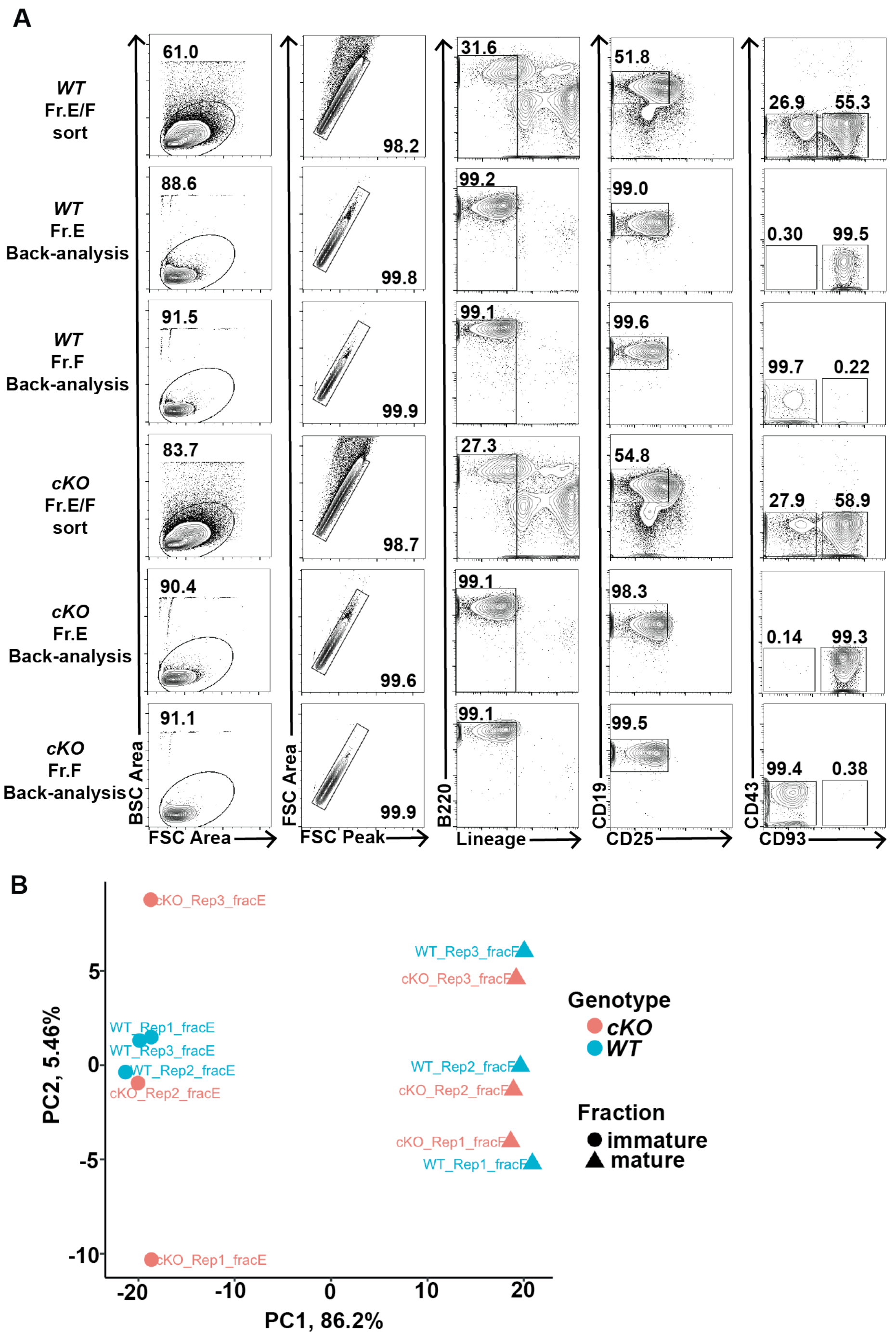


**Fig. S5. Purity of FACS sorted immature and mature bone marrow B cells and relative variation between RNA-seq replicates.** (**A**) Representative flow cytometric analyses of purified immature (Fr.E) and mature (Fr.F) bone marrow B cells from *WT* and *Srpk3-cKO* male mice for downstream RNA-seq analysis. (**B**) Principal component (PC) analysis of biological replicates from RNA-seq experiment. All data are representative of at least three independent experiments.

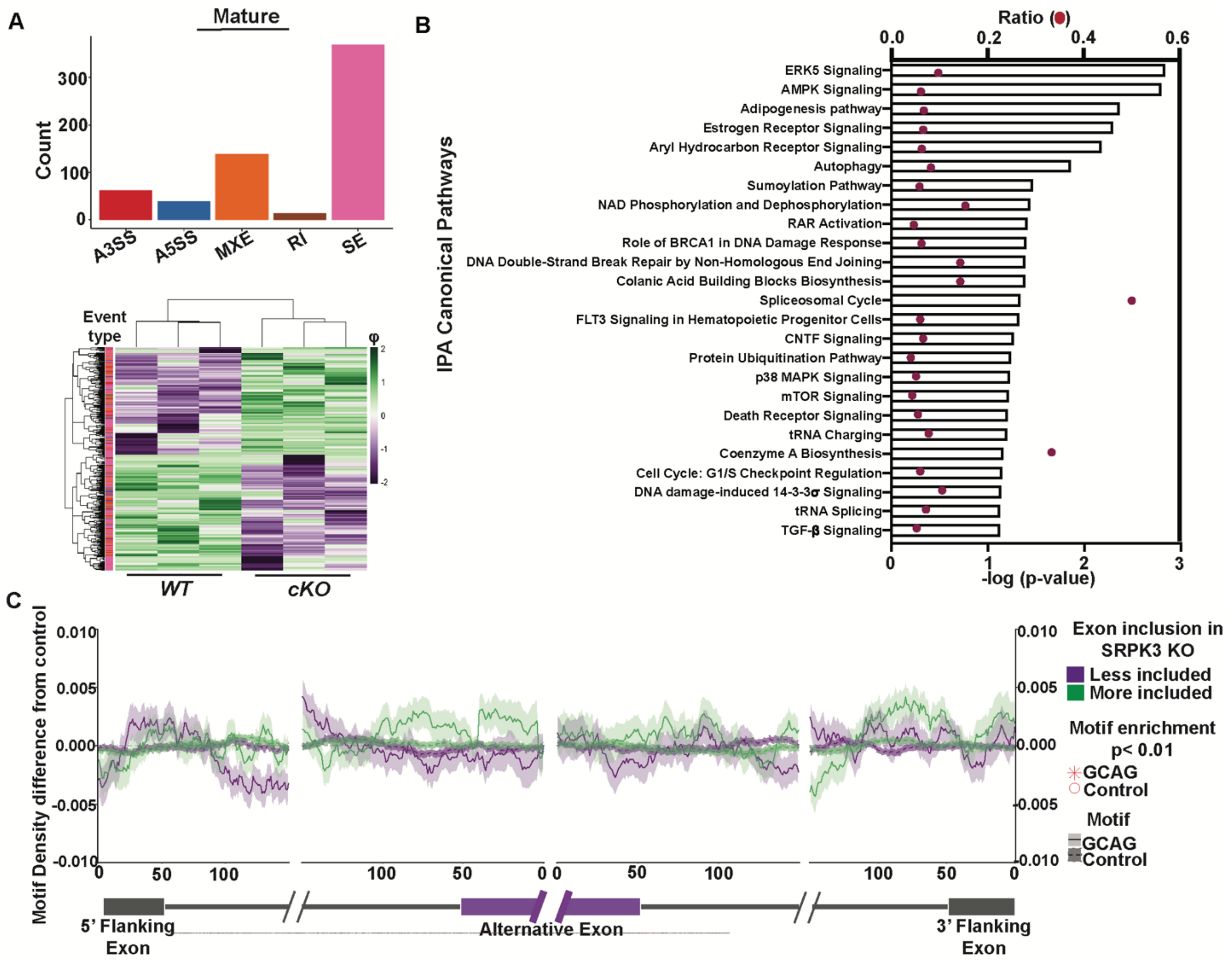


**Fig. S6. SRPK3 regulates alternative splicing patterns in mature bone marrow B cells.** (**A**) Significantly altered splicing event types in purified mature bone marrow B cells between *WT* (n=3) and *Srpk3*-*cKO* (n=3) male mice. AS event frequencies are displayed for alternative 3' splice sites (A3SS), alternative 5' splice sites (A5SS), mutually exclusive exons (MXE), retained introns (RI), and skipped exons (SE). Heatmaps show ψ values. Color (purple to green) represents z-score of ψ values across rows. $\Delta\psi \geq 0.1$, $FDR < 0.05$. No overlap in ψ between replicates across conditions. (**B**) IPA enrichment for the top 25 most significantly enriched canonical pathways in the mature bone marrow B cell RNA-seq splicing dataset. (**C**) Motif density surrounding SPRK3-sensitive exons. GCAG motif densities were calculated in the sequences surrounding exons that were more included or less included in the *Srpk3-cKO* relative to *WT*. Densities were also calculated for unaffected, control exons. The difference in densities between affected and control exons is plotted on the y-axis, with the shaded area representing the standard deviation of the difference. As a further control, the average motif density for a set of control motifs was also calculated (dotted line). These motifs were all 4-mers that contained the same GC and CpG content as GCAG. P-values for the enrichments were calculated using a Mann-Whitney U test comparing densities in affected and control exons. Positions were signified as significantly enriched if $P<0.01$.

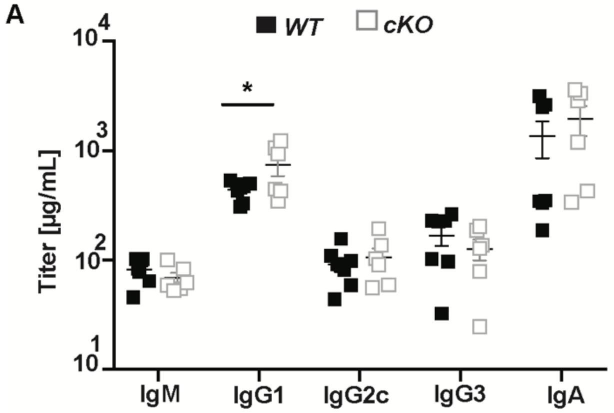


**Fig. S7. Elevated IgG1 immunoglobulin levels in *Srpk3-cKO* naïve mice.** (**A**) ELISA detection of immunoglobulin titers from mouse serum from naïve *WT* (n=7) and *Srpk3-cKO* (n=6) male mice. Two-tailed, Student's t-test was performed to determine statistical significance. *$P<0.05$. Graphs represent arithmetic mean ± SEM. All data are representative of at least three independent experiments.

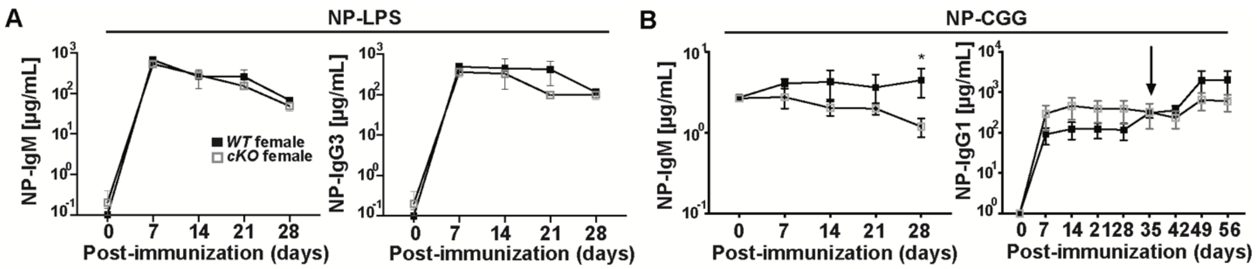


**Fig. S8. T lymphocyte-independent type-1 and T lymphocyte-dependent humoral immune responses in *Srpk3-cKO* mice.** (**A**) NP-specific ELISA detection of IgM and IgG3 immunoglobulin titers post-immunization with NP-LPS in female *WT* (n=2) and *Srpk3-cKO* (n=2) mice. (**B**) NP-specific ELISA detection of IgM and IgG1 immunoglobulin titers post-immunization with NP-CGG in female *WT* (n=3-4) and *Srpk3-cKO* (n=3-4) mice. Arrow indicates secondary boost (day 28) with respective antigen. 2-way ANOVA was performed to determine statistical significance. *P<0.05. Graphs represent arithmetic mean ± SEM.

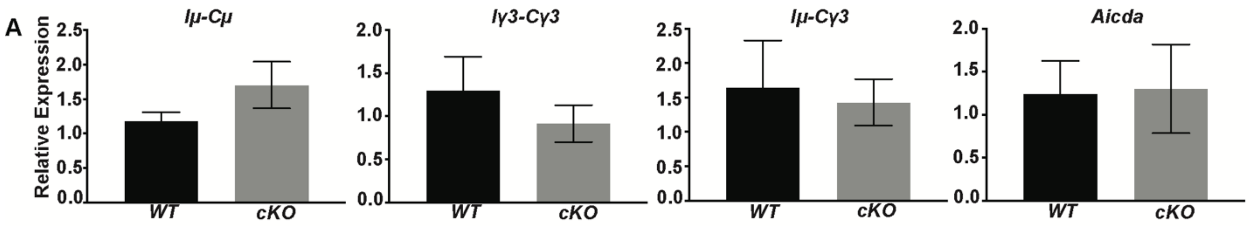


**Fig. S9. NP-Ficoll immunized mice are able to undergo class switch recombination.** (**A**) Relative expression of germline, post-switch and *AID* transcripts from purified splenic B cells harvested from NP-Ficoll immunized *WT* (n=3) and *Srpk3-cKO* (n=3) male mice 4 days post-immunization. Two-tailed, Student's t-test was performed to determine statistical significance. Graphs represent arithmetic mean ± SEM.

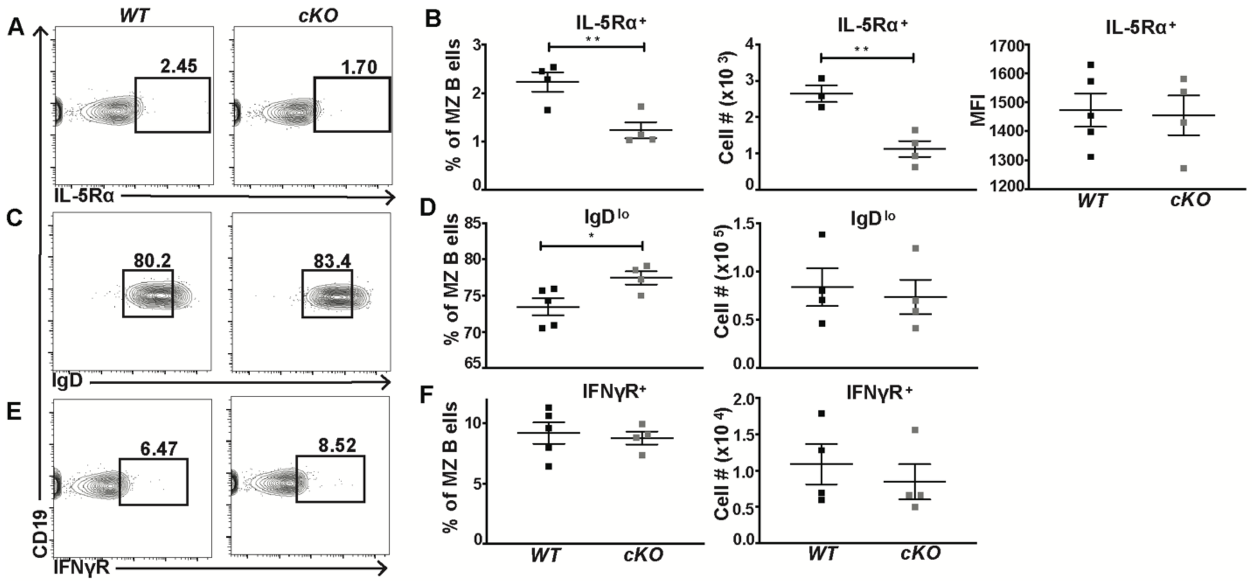


**Fig. S10. Decreased frequencies of IL-5Rα expressing and elevated frequencies of IgD expressing marginal zone B cells in *Srpk3-cKO* naïve mice.** (**A**) Flow cytometric analyses and (**B**) quantification and MFI of IL-5Rα$^+$ marginal zone B cells in *WT* (n=3-5) and *Srpk3-cKO* (n=4) male mice. (**C**) Flow cytometry and (**D**) quantification of IgD$^+$ marginal zone B cells in *WT* (n=4-5) and *Srpk3-cKO* (n=4) male mice. (**E**) Flow cytometry and (**F**) quantification of IFNγR$^+$ marginal zone B cells in *WT* (n=4-5) and *Srpk3-cKO* (n=4) male mice. Asterisks indicate statistical significance compared to *WT* littermate controls using either two-tailed, Student's t-test. *P<0.05, **P<0.005. Graphs represent arithmetic mean ± SEM. All data are representative of at least three independent experiments.

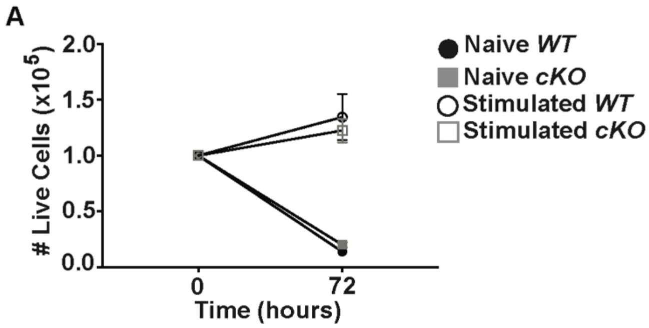


**Fig. S11. *Srpk3-cKO* marginal zone B cells are capable of proliferating in response to stimulation in vitro.** (**A**) Live cell counts of unstimulated (naïve) or stimulated *WT* (n=3) and *Srpk3-cKO* (n=3) marginal zone B cells 3 days post-stimulation with α-δ-dex, IL-5 and IFNg. 2-way ANOVA was performed to determine statistical significance. Graphs represent arithmetic mean ± SEM. All data are representative of at least three independent experiments.

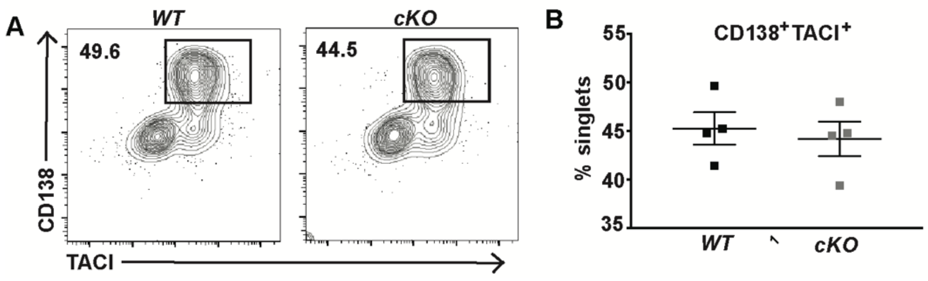


**Fig. S12. *Srpk3-cKO* marginal zone B cells differentiate into plasma cells in vitro.** (**A**) Flow cytometric analysis of CD138 and TACI and (**B**) frequency of dual CD138- and TACI-expressing cells after 3 days of in vitro stimulation with α-δ-dex, IL-5 and IFNg of purified marginal zone B cells from *WT* (n=4) and *Srpk3-cKO* (n=4) male mice. Two-tailed, Student's t-test was performed to determine statistical significance. Graphs represent arithmetic mean ± SEM. All data are representative of at least three independent experiments.

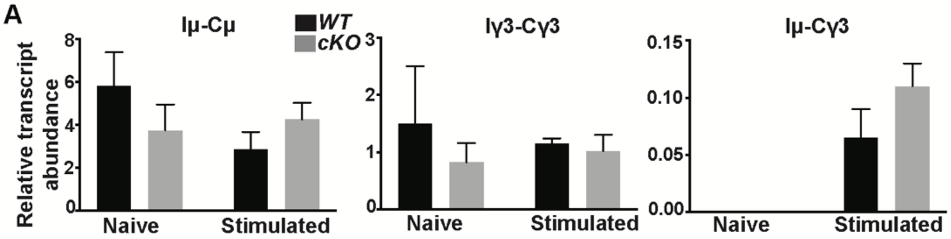


**Fig. S13. In vitro stimulated *Srpk3-cKO* marginal zone B cells can undergo class switch recombination.** (**A**) Relative expression of germline and post-switch transcripts from naïve or stimulated (α-δ-dex, IL-5 and IFNg) marginal zone B cells isolated from *WT* (n=2) and *Srpk3-cKO* (n=2) male mice. 2-way ANOVA was performed to determine statistical significance. Graphs represent arithmetic mean ± SEM.

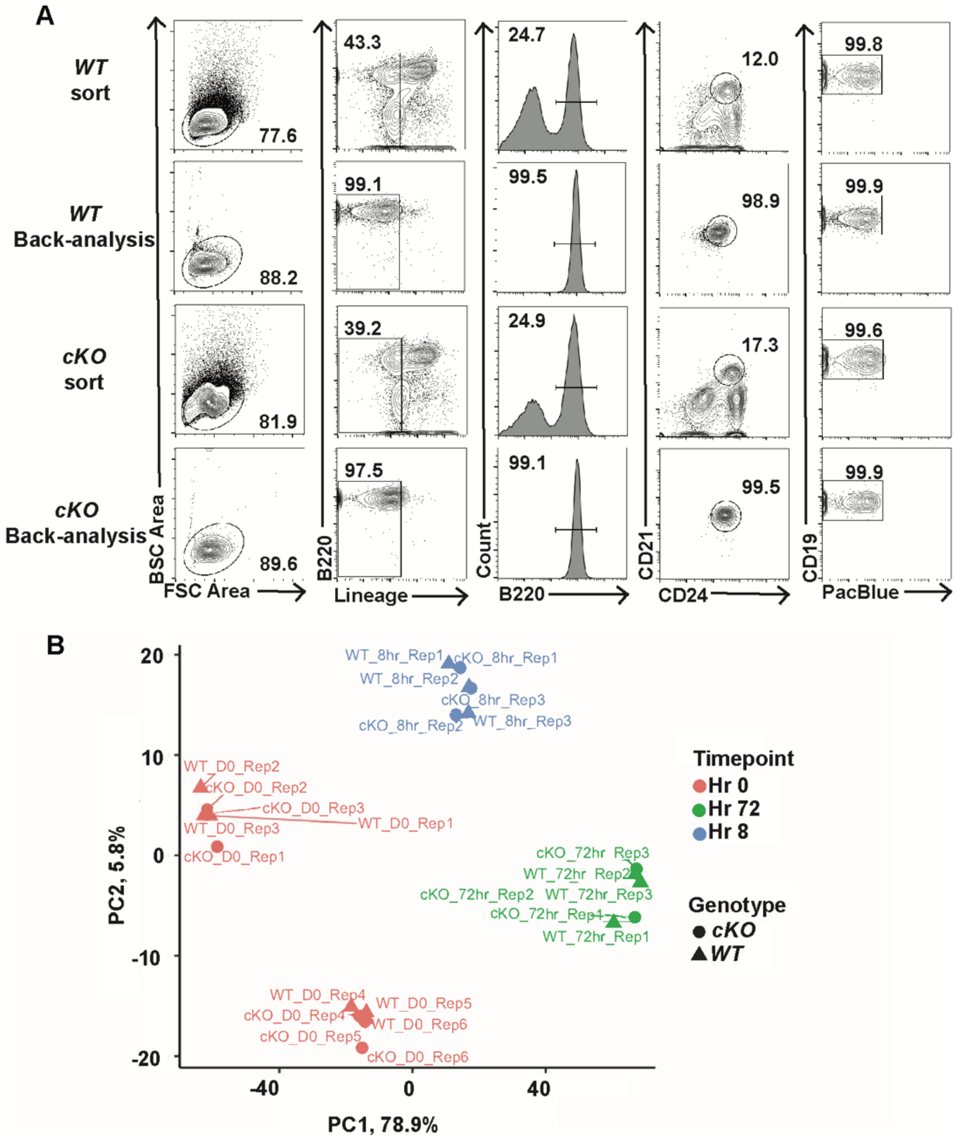


**Fig. S14. Purity of FACS sorted marginal zone B cells and relative variation between RNA-seq replicates across conditions.** (**A**) Representative flow data of purified marginal zone B cells from *WT* and *Srpk3-cKO* male mice for downstream RNA-seq analysis. (**B**) PC analysis of biological replicates from RNA-seq experiment across conditions. All data are representative of at least three independent experiments.

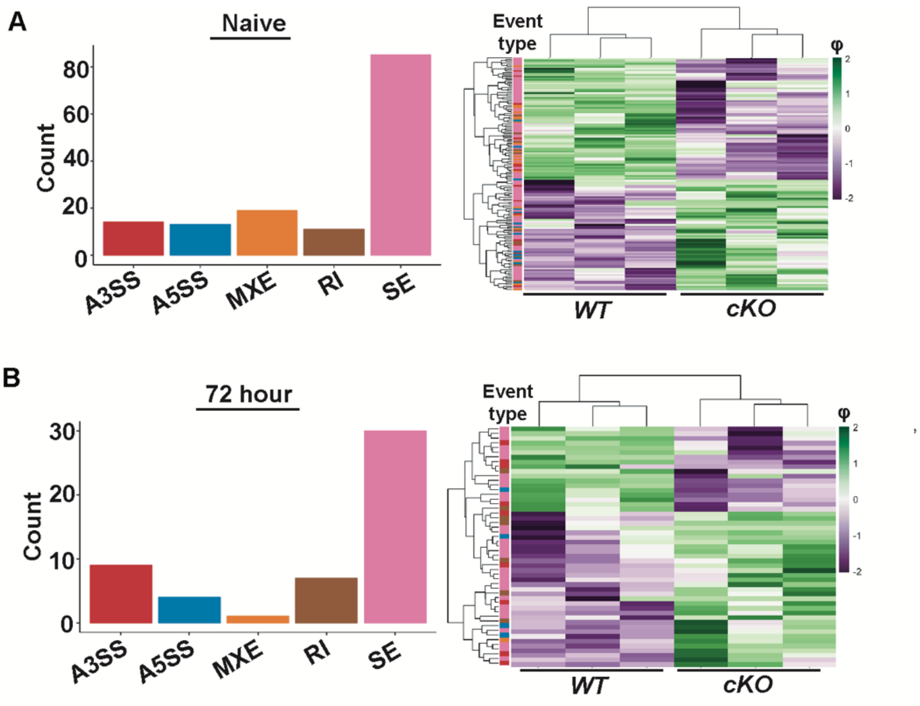


**Fig. S15. SRPK3 is necessary for regulating alternative splicing patterns in naïve and 72-hour stimulated marginal zone B cells.** (**A**) Significantly altered splicing event types in naïve marginal zone B cells purified from *WT* (n=3) and *Srpk3-cKO* (n=3) male mice. (**B**) Significantly altered splicing event types in 72-hour stimulated marginal zone B cells purified from *WT* (n=3) and *Srpk3-cKO* (n=3) male mice. Same mice were used to split purified marginal zone B cells between conditions. Heatmaps show ψ values. Color represents z-score of ψ values across rows. $\Delta\psi \geq 0.1$, FDR < 0.05. No overlap in ψ between replicates across conditions.

Table S1. PCR Primer List

| | |
|---|---|
| **Genotype PCR & Gene deletion** | |
| *Srpk3*-R | TCTGTCTCACCCTTATTCCCAACCC |
| *Srpk3*-F | GCTGACTCTGCCAAGTAAAAGGACC |
| *Srpk3*-ttR | GAGTGAAGGGCAGTCAACAACAC |
| *Cd79a-Cre-F* | ACACACCTGGAAGATGCTCCTGTC |
| *Cd79a-Cre-R* | CAAAGTCAGTGCGTTCAAAGGC |
| *Hprt-F* | GGGGGCTATAAGTTCTTTGCTGACC |
| *Hprt-R* | CCTGTATCCAACACTTCGAGAGGTCC |
| *Srpk3-F* | CCCTGAAAGTGGTGAAGAGTG |
| *Srpk3-R* | GCAGACCCTGGTAGTTGGATTTG |
| **Class switch recombination** | |
| *Iμ-Cγ3-F* | CTCGGTGGCTTTGAAGGAAC |
| *Iμ-Cγ3-R* | ACCAAGGGATAGACAGATGGGG |
| *Iμ-Cμ-F* | ACCTGGGAATGTATGGTTGTGGCTT |
| *Iμ-Cμ-R* | TCTGAACCTTCAAGGATGCTCTTG |
| *Iγ3-Cγ3-F* | AACTACTGCTACCACCACCACCAG |
| *Iγ3-Cγ3-R* | ACCAAGGGATAGACAGATGGGG |
| *Aicda-F* | TGCTACGTGGTGAAGAGGAG |
| *Aicda-R* | TCCCAGTCTGAGATGTAGCG |
| **RNAseq validation** | |
| *Slc1a5-F1* | CATCTCCGCTCTTCTCTTCCC |
| *Slc1a5-R1* | CTGAGCTCGGCATCTTGGTT |
| *Slc1a5-F2* | CACCATCCTGGTCACAGCCA |
| *Slc1a5-R2* | CTGAGCTCGGCATCTTGGTT |
| *Mief2-F1* | AATGCTCGATTGGTGCTAGGT |
| *Mief2-R1* | CCCCTTGGTGTCATCCTCAT |
| *Mief2-F2* | AAAGCGGTGGAGCCCACG |
| *Mief2-R2* | AACTTAGGTGTCGTGGGAGAA |
| *Mief1-F* | TTGACACAGGAGAAACTTCTT |
| *Mief1-R* | ATGGTCAGCCGTCACCAC |
| *Nnt-F1* | GCACAGCTCTGATTCCAG |
| *Nnt-R1* | CGATCGTCAGGCTTCTTCGA |
| *Nnt-F2* | GATTCCAGATCATGTACCT |
| *Nnt-R2* | ATCTGGATCCGCTTTGCGAT |
| *Nek4-F1* | GATCAAGTTACTGGTATTATAG |
| *Nek4-R1* | ACCCAGCCACAGGTCCACC |
| *Nek4-F2* | GCCAGACCAGTTCCAAGAATTAC |
| *Nek4-R2* | GCCAGAGAGAGGAGACGGC |
| *Tyk2-F1* | AGAGACTTCCTCAGACCGGG |
| *Tyk2-R1* | GGTGTGGGAGCTTTTTGCTC |
| *Tyk2-F2* | CGAATGAGGGATGGTATGTTG |
| *Tyk2-R2* | GCTGCCGGATGTGCTGTC |
